# Cultural affiliation accounts for most of the spatiotemporal variation in burial rite practices

**DOI:** 10.64898/2026.05.25.725982

**Authors:** Elisabetta Canteri, Robert Staniuk, Adrian Timpson, Peter Schauer, Jelena Bulatović, Maria Ivanova-Bieg, Samantha S. Reiter, Helene Agerskov Rose, Jan Kolář, Mark G. Thomas, Fernando Racimo, Stephen Shennan

## Abstract

Describing and interpreting spatiotemporal patterns in human culture has been a central focus of anthropology and archaeology for over a century. Recent ethnographic studies have highlighted the complexity of the processes generating these patterns, including isolation-by-distance, homophily, and common descent. However, investigating these processes in prehistoric archaeology remains challenging. Here we make use of a new interdisciplinary database and a combined dataset of ancient DNA (aDNA) genomic sequences to analyse the relationship between spatiotemporal patterns in cultural and genomic variation, by testing whether broadly defined clusters of genomic affinities correspond to spatiotemporal changes in burial rites, while controlling for other factors, using a Gaussian process model. We use data from the Big Interdisciplinary Archaeological Database (BIAD), linking mortuary information from ∼4,200 individuals with genetic ancestry and mobility data inferred from over 1,300 human genomes, from Western Eurasia ∼10,000-2000 BP. By integrating and modelling these diverse datasets, we aim to provide a detailed understanding of how genomic history intersects with cultural evolution, offering new insights into the dynamics behind these complex processes, and the extent to which genes and culture are transmitted in parallel. In the case of burial orientation, we found that cultural affiliation was the main factor accounting for variation with little to no role for ancestry, while for body position the picture was more mixed but cultural affiliation also played an important role.

## Introduction

Characterising the ways in which human culture varies over space and time and explaining the causes behind this variation have been long-standing goals of archaeology and anthropology (e.g., Boas, 1896). Early in the 20th century, Clark Wissler’s (1923) anthropological ‘culture area’ approach grouped together North American indigenous societies based on key cultural traits. Similarly, in archaeology, the concept of ‘culture history’ emerged around the same time: the idea that history is composed of successive ‘cultures’, discrete complexes of associated material culture traits that appear, have a period of currency and then disappear, to be replaced by others. These were regarded as the material expression of what today we would call “a people” (Childe, 1929, v-vi). Subsequently, archaeological cultures came to be regarded as purely typological entities based on material culture, in response to the early 20th century association between ‘cultures’ and ‘races’.

In more recent years, there has been a renewed interest in the processes that generate such patterns, following on the emergence of the new discipline of cultural evolution (e.g., Mesoudi, 2011). The most basic mechanism that can generate spatial patterns is ‘isolation-by distance’: as geographic distance between people increases, they will be less likely to interact with one another, reproduce together (in genetics, see for example, Novembre et al., 2008; Slatkin, 1993; Wright, 1943), and thus less likely to transmit and share cultural traits. Similarly, as distance in time between people increases, there is greater opportunity for change to occur simply due to stochastic variation and evolution in cultural and genetic traits during the process of transmission (Loog et al., 2017).

A key question then is whether there are further processes at work generating boundaries in space and time not accounted for by isolation-by distance or evolution. One possibility is differences reflecting adaptations to spatiotemporally heterogeneous environments. Another well-known one is homophily: the preference for interaction with people like oneself in some way. In populations that are initially well-mixed even a slight preference for people with some trait can rapidly generate population structure dividing those with the trait from those without (Axelrod, 1997).

Several anthropological studies have explored the importance of isolation-by-distance compared to homophily or environmental factors. For example, Ross et al. (2013) found that geographical distance and ethnolinguistic affiliation had significant independent effects on diversity in folk tales: ‘folktales from the same culture [ethnolinguistic population] found 100 km apart are, on average, as similar as folktales found 10 km apart in different cultures (see also Barbieri et al., 2022). However, shared language can not only increase interaction, it also implies a shared transmission history, so the two are likely to reinforce one another in generating boundaries. In another study, Mathew and Perreault (2015) used data from the Western North American Indian database (Jorgensen, 1980) to test whether trait variation between ethnolinguistic groups in a large number of different domains (including technology, subsistence, marriage, kinship and ceremony) could be accounted for by ecological factors, spatial distance or cultural history, represented by distances between the groups on a linguistic tree. Cultural history had a greater effect than ecology in the great majority of cases and the effect of cultural phylogeny was much more important than spatial distance; in other words, within-group transmission was much more important than between-group diffusion over significant periods of time. An ethnohistorical study of multiple material culture traditions in California hunter-gatherer societies (Jordan & Shennan, 2009) found that three of them – basketry, cradles and ceremonial dress – showed similar patterns of branching descent linked to language differences, while a fourth—earth lodges—had an entirely distinct transmission history. This points to the existence of distinct ‘cultural packages’ with different histories within these societies (Boyd et al., 1997).

Compared to anthropology, the number of archaeological studies addressing the importance of isolation-by-distance in the structuring of cultural differences and similarities is much more limited. Cross-cultural data sets are much less common in this field. For the majority of prehistory, we have no knowledge of the languages spoken or of the definition of ethnolinguistic groups, or at least not enough to structure an analysis of this nature. Based on a set of descriptive attributes for Neolithic ceramic and personal ornament assemblages in Europe, Shennan et al. (2015) tested whether isolation-by-distance in space and/or time was sufficient to account for cultural space-time patterning, or whether ‘cultures’ represented in some sense ‘real entities’ imposing boundaries on patterning that reflected preferential interaction and/or shared descent. Both sets of data showed significant population structure related to cultural affiliation, much stronger in the case of pottery than ornaments, which might be expected given that European Neolithic cultures are generally defined on the basis of ceramics. The study showed that it was possible to identify population structuring beyond isolation-by-distance based on archaeological evidence ‘without continuing to attempt the fruitless task of correlating its patterns with past ethnolinguistic units’ (Shennan et al., 2015).

In recent years, discussions about the nature of archaeologists’ culture-historical entities and their significance have been revived by the results of whole genome ancient DNA (aDNA) studies of prehistoric populations, especially in Europe (e.g., Allentoft et al., 2015; Haak et al., 2015; Hofmanova et al., 2016; Lazaridis et al., 2016). These have been controversial because they seemed to support the early culture history assumption that archaeological cultures correspond to particular biological populations (Frieman & Hofmann, 2019; Furholt, 2017; Kristiansen, 2022). These discussions have generally failed to embrace the kinds of cultural evolutionary studies (described above) that have addressed transmission processes. aDNA studies offer the possibility of relating cultural patterns to genetic descent and evaluating the correlation(s) between them. Thus, we can avoid the simplistic opposition between culture and genes that characterises most of the debates, making them so acrimonious. Yet, the analysis of population level variation in cultural traits in relation to genomic variation is faced with a significant problem: archaeological cultural data are routinely described in a culture-specific way. As a result, the data are generally described in terms of essentialist types rather than constellations of attributes (for exceptions see Furholt, 2009; Shennan et al., 2015). Culturally-charged terms like ‘Corded Ware Beakers’ and ‘Bell Beakers’ obscure variation in material culture, which creates additional challenges for the evaluation of continuous variation in space and time (Furholt, 2019). If our aim is to re-evaluate the role of archaeological cultures, a wholesale redescription of the cultural archaeological record is needed (Shennan et al., 2015).

However, some cultural features of the archaeological record are often described in terms that are agnostic to previously defined cultural terms. This is the case with burial practices, which have long been of archaeological interest because they relate to an important aspect of human lives that is governed by social norms concerning what is appropriate (Fahlander & Oestigaard, 2008; Gramsch, 2013; Nilsson Stutz, 2003). Burial practices themselves comprise different domains that may not be subject to the same considerations. For example, norms affecting what grave goods are appropriate may differ from those affecting the treatment of the body (Carr, 1995, table XI).

In this paper we will focus on two important aspects of inhumation burials: the way the buried individual is placed in the grave and their orientation in relation to the points of the compass. These have been the subject of several studies that explored how variation in these domains relates to different archaeological cultures. Häusler’s (2001) pioneering, large-scale investigation into the patterning of body positioning and orientation in Western Eurasia between the Neolithic and Bronze Age provided evidence of the complex patterning of funerary traits through time and space. However, for every norm he documented, a preceding or following modification of the practice was documented, leading him to conclude that no underlying rule could explain the logic of burials beyond religious or magical practices. Carr’s (1995) ethnographic survey of the determinants of different aspects of burial variation found that body position and orientation were most influenced by beliefs about universal orders and the afterlife, but associations were not very strong.

Here, using spatiotemporal modelling under Bayesian inference (Gómez-Rubio, 2020), we test whether variation in these domains is simply determined by isolation-by-distance in space and time; whether it relates to the genomic affinities of the individual concerned (implying that burial norms may have at least partly been transmitted vertically, in parallel with the genes); or whether they are more strongly related to differences between archaeological cultures (implying that these do have a bearing on the social networks through which burial norms were transmitted).

## Materials and Methods

### Data

#### Grave individuals

Data on grave individuals was collected as part of the ERC-funded COREX project from published archaeological research (monographs, articles, databases) and included in the Big Interdisciplinary Archaeological Database (BIAD; https://biadwiki.org/) being built by the project. The protocol for data collection followed a hierarchical data structure of increasing granularity from spatial data on the location of the discovered individual (Table: Sites), contemporary temporal aggregations of finds (Table: Phases), find circumstances of the individual (Table: Graves), demographic attributes of the individual (Table: Individuals), details of burial rite (Table: Rites), and accompanying material culture (Table: MaterialCulture). Absolute dating such as radiocarbon dating was collected as an independent variable primarily associated with the spatiotemporal positioning of the date (Table: C14Samples), which could be directly linked to an individual via horizontal connection (Table: Items). The data was uploaded to the MySQL database (BIAD) with the set of primary keys (e.g., SiteID, PhaseID, GraveID, IndividualID, RiteID, aDNAID) used for querying. Unlike primary keys linked to archaeological data, the information allowing direct association of aDNA data was made possible through the implementation of aDNAID, which was associated with the specific individual. This unique identifier, or identifiers in case of re-sampling, was used to query an existing aDNA repository (AADR: the Allen Ancient DNA Resource), for matching genomic data.

#### Data collection protocol

Site name and, if possible, coordinates were obtained directly from the published sources. If no coordinates were provided, an attempt at identifying site location was made using Google Earth Pro to first identify the nearest possible matching toponym and, depending on the data available in the resource, obtain the closest possible coordinates to the original place of discovery. Due to the heterogeneity of the data sources considered in this study, which include sites discovered in the late 19th century and originating from various locations across Western Eurasia, this was not always possible. Where attempts at precise identification were impossible, a point estimate to the nearest matching toponym and possible extent of spatial error was provided (usually measuring ca. 5 km radius).

Archaeological phasing of the discoveries made at the location followed that reported by the primary investigators as much as possible, unless modified findings were reported by later researchers incorporating dating methods substantially modifying the initial findings. In cases where multiple strands of data were encountered (e.g., archaeobotanical, archaeozoological, human palaeodemographic data), we considered it a priority to maintain or enable interoperability between the different types of information, rather than preserving high-resolution but unsynchronized data. To that end, aggregations resulting in decreasing resolutions were proposed and discussed between the data collectors. In cases where no phase information was provided, especially if no references to findings from preceding or following phases was stated, reported findings were used to propose a phase and aggregate information. Information about find circumstances were extracted directly from the published resource, and where no detailed data was encountered, attempts were made to obtain the original publication. Key information was grave depth, grave construction and the minimal number of individuals (MNI) in the grave.

Data on individuals were extracted directly from published works, and similarly to graves, if no detailed data was found, attempts at acquiring this information from other published sources were made. The key variables were demographic data (categorical and continuous anthropological age and sex) and unique identifiers for genetic sequences.

Archaeological information was collected for a total of 4,275 individuals, including: country of origin, site name and geographic coordinates, archaeological period, phase, culture, calibrated radiocarbon date and error, deposition type, body positioning, burial side, minimum and maximum burial orientation measured in angle degrees from north (0°), and presence of grave goods and ochre.

**Table 1.** Burial practices. Burial practices covered in this study, with a short description of the traits and variables considered and how the data was collected.

| Trait | Description | Data collection | Variables |
| --- | --- | --- | --- |
| *Burial position* | The overall position of the deceased | Recorded directly from sources, converted to one of the general types, visual data inspected if available | Crouched<br>Extended<br>Sitting<br>Inside a vessel |
| *Burial side* | Rotation axis of the deceased | Recorded directly from sources, converted to one of the general types, visual data inspected if available | Left<br>Right<br>Back<br>Front<br>Bottom |
| *Burial orientation* | Arrangement of the body according to the cardinal directions | Orientation recorded, converted into angular data with confidence intervals adjusted for the accuracy of reporting | Min-max between 0-360° |

#### ADMIXTURE proportions and mobility estimates from human aDNA studies

Estimates of ancestry clustering proportions as inferred by the software ADMIXTURE (Alexander et al., 2009) on a set of more than 1,600 imputed ancient genomes from across Eurasia were accessed from Allentoft et al. (2024). These correspond to a broad-scale clustering analysis, and we specifically focused on four ancestries that are maximized among, respectively, Neolithic farmers migrating from Anatolia (NEOL), Western hunter-gatherers (WHG), Eastern hunter-gatherers (EHG) and Caucasus hunter-gatherers (CHG), which are generally agreed to represent various streams of ancestry that contributed to the bulk of the genetic make-up of Western Eurasian peoples over the course of the Neolithic and Bronze Age (Allentoft et al., 2015; Haak et al., 2015; Hofmanova et al., 2016; Lazaridis et al., 2016). Even though we henceforth call these “ancestry values”, we note that these estimates are merely the output of a specific soft clustering algorithm, which assigns fractional cluster memberships to all individuals on which it is applied. These clusters do not necessarily correspond to any specific group of people that lived at a particular time and place; they are only an approximation to the true history of the genomes under study. Specifically, the ancestry estimates represent a rough and incomplete summary of a much more complex set of genetic relationships over space and time, i.e. the full ancestral recombination graph.

Genetically-inferred mobility estimates were calculated using the methods in Schmid and Schiffels (2023), which analysed genomic sequences from the Allen Ancient DNA Resource (AADR) V50.0 (David Reich Lab, 2021). We decided to calculate mobility estimates using retrospection distances based on a temporal kernel size of 250 years, as this would roughly correspond to the temporal resolution of our data; thus, we estimated the spatial location of maximum genetic similarity to the buried individual and the resulting “genetic relocation” at 250 years before that individual.

To pair the genetic estimates of ancestry and mobility with the grave individuals (Figure 1), we first checked for matches among the three datasets using the aDNA primary key (“aDNAID”). In case of a positive match (ancestry individuals N = 144, mobility individuals N = 462), meaning that the genetic information belongs to the specific grave individual, we assigned the corresponding estimates of ancestry and mobility. For all other grave individuals, we assigned the average value across up to the four closest genomes within a distance radius of 350 km and 300 years.

**Figure 1.**
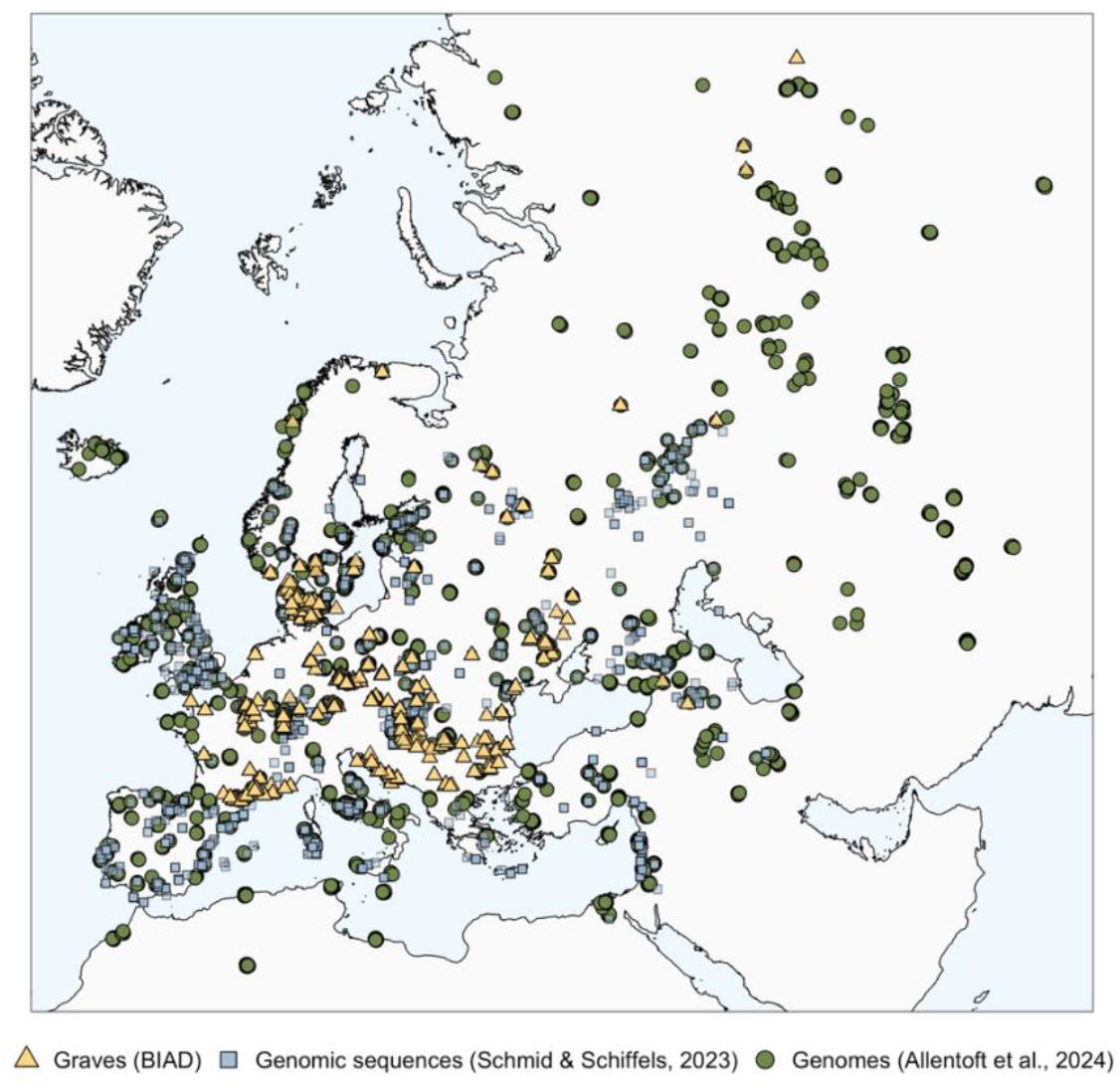
Geographic distribution of the data. Locations of the grave individuals (yellow triangles), the ancient genomic sequences analysed in Schmid and Schiffels (2023) used to infer mobility (blue squares), and the ancient genomes from Allentoft et al. (2024) used for estimating ancestry proportions (green circles).

After removing individuals outside the date range of our interest (> 11,000 calBP) and individuals without date information, the final dataset includes a total of 3,995 individuals, of which 3,210 have genetic ancestry information and 3,155 have genetically inferred mobility estimates. Individuals span a temporal interval between 10,600 calBP and 760 calBP.

#### Data exploration

We first explored patterns related to the side on which an individual was buried (e.g., left, back, right, front) and body positionings (e.g., crouched, extended, sitting, inside a vessel) (Figure 2, Figure S1). Our dataset includes 1,515 individuals with burial side information and 2,584 individuals with information on body positioning.

**Figure 2.**
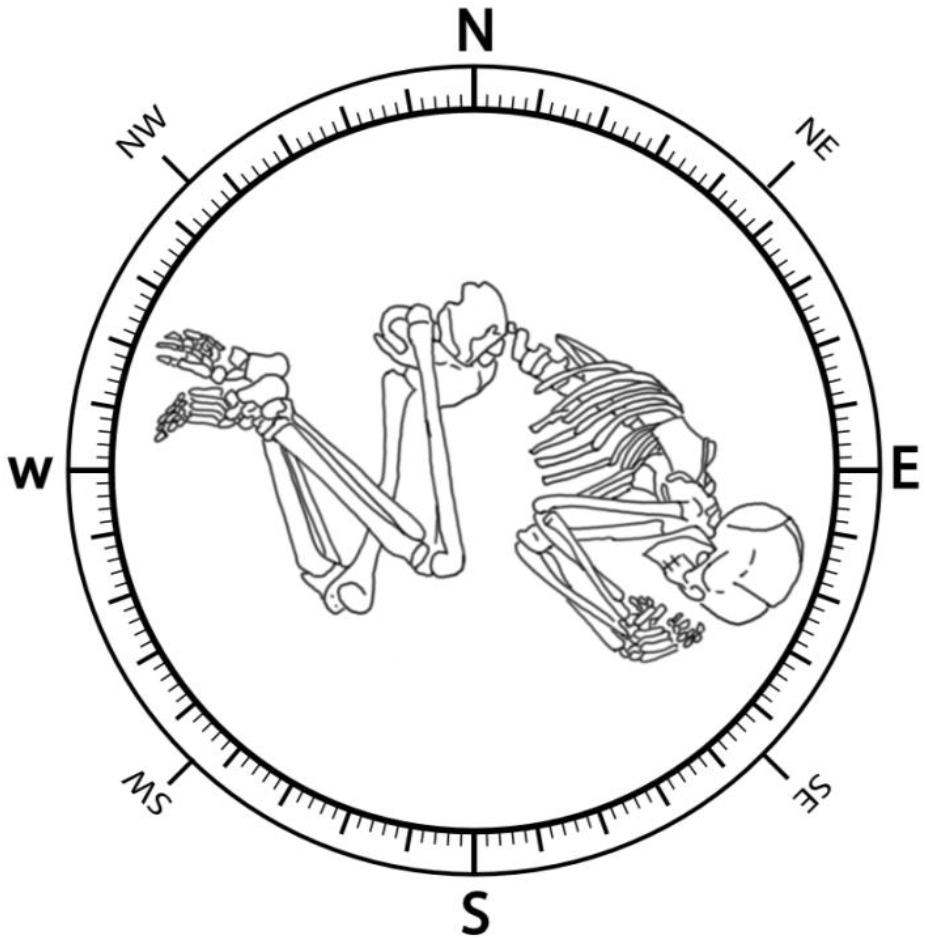
Body position, burial side, and burial orientation. Conceptual figure showing the traits considered in this study. The skeleton drawing has been adapted from Neugebauer-Maresch and Lenneis (2015). Compass designed by rawpixel.com / Freepik.

We used Multiple Correspondence Analysis (MCA), implemented using the ‘factoextra’ (Kassambara & Mundt, 2026) and ‘FactoMineR’ (Lê et al., 2008) packages in R, and UpSet plot, implemented using the ‘UpSetR’ R package (Conway et al., 2017), to identify intersections and structures in the data. We then explored how body-position and burial-side relate to time (archaeological period), culture (10 most represented cultures in our data with information on burial orientation), sex, genetic ancestry, and mobility. Cultures considered are: Linearbandkeramik (LBK), Lažňany, Sopot, Bell Beaker complex (hereby referred as Bell Beaker), Únětice, Corded Ware/Single Grave/Battle Axe/Fatyanovo (hereafter referred to as Corded Ware), Baden, Dnieper-Donets, Trichterbecher, Baltic Mesolithic-Neolithic (hereafter referred to as Baltic Meso-Neolithic).

### Modelling the spatiotemporal effects of ancestry, mobility, and culture on burial side

Based on the results of the MCA and UpSet plot (Figure 3), we decided to focus on the left, right and back burial sides. We used the Bayesian Latent Gaussian modelling approach implemented in INLA and the ‘inlabru’ R package (Bachl et al., 2019; Rue et al., 2009), to model the burial sides against values of genetic ancestry and mobility, and cultural association, via a spatiotemporal Gaussian process model.

**Figure 3.**
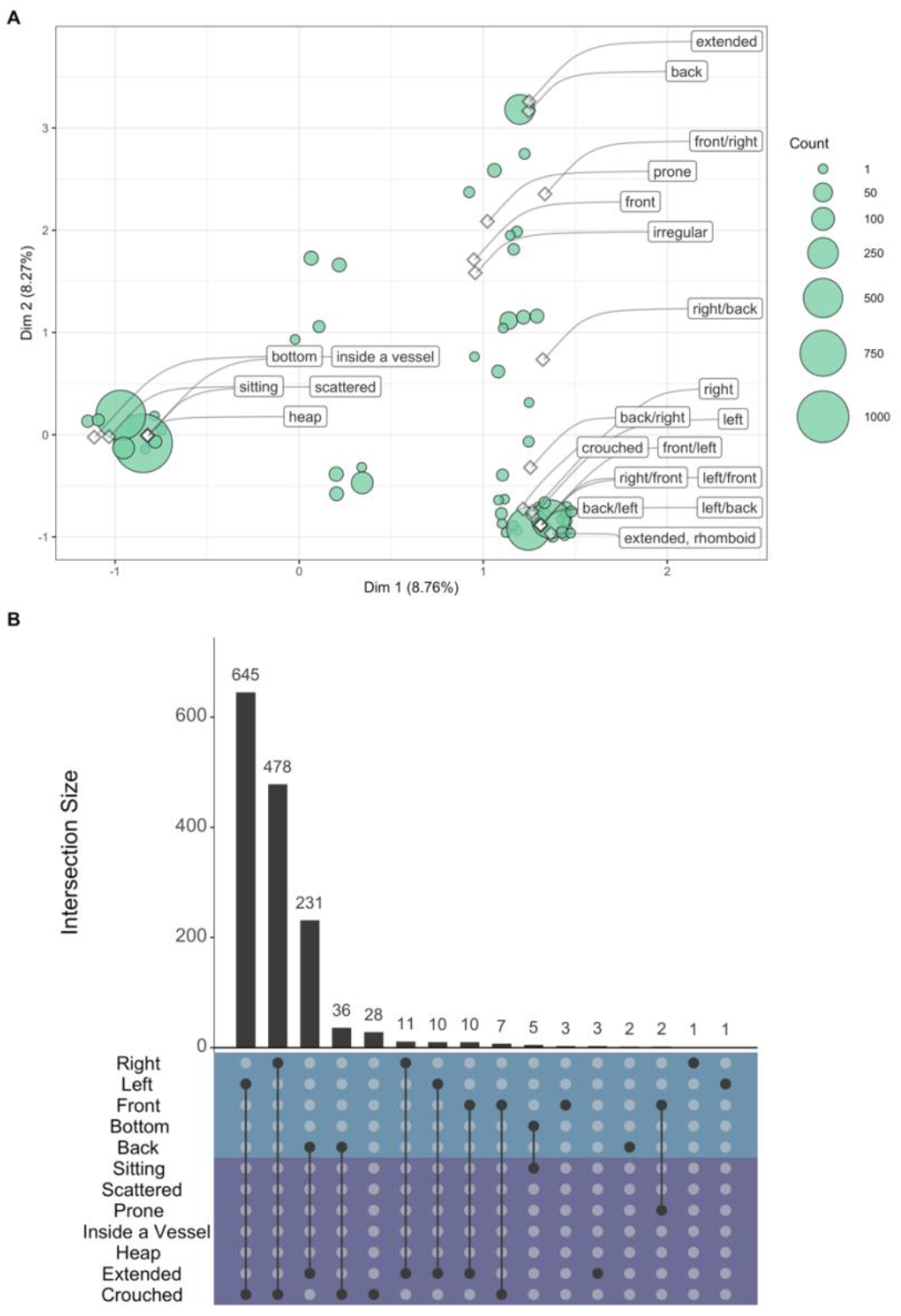
Structure in burial rite traditions. **A)** Multiple Correspondence Analysis (MCA) showing clustering in burial side and body positioning of grave individuals. The size of the circles represents the number of individuals having the same coordinates in the MCA space. Only the first two dimensions are plotted, and an overview of the variance explained by all dimensions is shown in Figure S5. **B)** UpSet plot showing intersection between burial sides and body positioning of grave individuals. Purple rows refer to body positioning while blue rows refer to burial side. A connection between two black dots indicates an intersection between variables. Black bars show the number of individuals showing that specific intersection. The number of individuals within each variable (set size) is shown in Figure S6.

In INLA, continuous-space models are based on a Stochastic Partial Differential Equation (SPDE) approach. This requires the construction of distinct sets of data and model objects, with specific priors. To take informed decisions about this prior information, we performed a set of preliminary spatial analyses. We started by creating a spatial mesh, which is used to provide a piecewise linear representation of the continuous spatial surface, based on a triangulation of the modelled region. We defined the spatial range (modelled region) by calculating the diagonal distance (Euclidean distance) between two vertices of the bounding box coordinates around the locations of the grave individuals. Based on fractions of the spatial range, we defined the maximum edge, cutoff, and offset parameters of the 2D spatial mesh for the Gaussian random field. We also created a 1D temporal mesh, with knots every 1,000 years between 0 and 12,000 cal BP.

The SPDE model object defines the model smoothness. It requires the specification of prior distributions for the variance and range parameters, and the mesh. We assume a Gaussian random field characterized by a Matérn covariance function with penalized complexity priors for the practical range and variation. To set the prior for the spatial range, we tested different models with different prior combinations and selected the one with the lowest Watanabe-Akaike Information Criterion (WAIC) score (Watanabe, 2010), which estimates out-of-sample predictive accuracy by integrating over the full posterior distribution, appropriately accounting for model complexity (Gelman et al., 2013). Unlike information criteria based on point estimates of the parameters, WAIC is valid for complex Bayesian models, including hierarchical models and models with informative priors, and is asymptotically equivalent to Bayesian leave-one-out cross-validation (Gelman et al., 2013). We first tried different bandwidths of a kernel density over the grave locations, to better understand spatial autocorrelation. We also tested for a homogeneous spatial point process (L-test). This is another way to test for spatial autocorrelation as clustered point patterns would deviate from a perfect homogeneous Poisson point process (i.e., a process where the larger the distance of the radius from a point, the more points you find). We used the “spatstat” R package (Baddeley et al., 2015) to estimate kernel densities with a default bandwidth, a likelihood cross validation bandwidth, a cross validated bandwidth and bandwidth values of 250, 200, 180 and 100. Based on the estimated densities, we tested SPDE models with values for the spatial range equal to the default bandwidth, the bandwidth selected using likelihood cross validation, the one selected using cross validation and the value resulting from the L-test. The model with the lowest WAIC score had the bandwidth selected with cross validation. We therefore used a SPDE model with spatial range set so that the probability of a range exceeding 98 km is 0.01. The prior for the variance explained by the spatial effect is set such that the probability of a standard deviation exceeding 1 is 0.01. The SPDE model represents the spatiotemporal process, and it is used as an explanatory variable when modelling burial-side.

For each burial side (left, right, back), we fit the following sets of models:

1. *Baseline: Burial Side ∼ spatiotemporal process*
2. *Baseline + mobility + ancestry (all ancestries)*
3. *Baseline + Eastern Hunter Gatherer (EHG) ancestry*
4. *Baseline + Western Hunter Gatherer (WHG) ancestry*
5. *Baseline + Caucasus Hunter Gatherer (CHG) ancestry*
6. *Baseline + Neolithic farmers (NEOL) ancestry*
7. *Baseline + mobility*

Due to the fractional nature of the ancestry estimates from ADMIXTURE, we found that there was strong collinearity between WHG and all the other ancestries (Figure S2), so we excluded the WHG fractions from any model in which the other ancestry values were also used (e.g. Model 2). All models were applied to data covering a period between ∼10,000 and ∼2,000 calBP.

We also ran a set of models before and after the arrival of the Steppe ancestry in Western Europe (∼ 5,000 calBP) (Haak et al., 2015). The models before 5,000 calBP are:

1. *Baseline*
2. *Baseline + ancestry (EHG, CHG, NEOL)*
3. *Baseline + ancestry + mobility*

The models after 5,000 calBP are:

1. *Baseline*
2. *Baseline + Steppe ancestry (EHG + CHG) + NEOL ancestry + mobility*
3. *Baseline + Steppe ancestry + NEOL ancestry*
4. *Baseline + Steppe ancestry*

While we observed the full posterior distribution of all models, we specifically focused on the 95% credible intervals of posterior densities of the coefficients for the covariates, to determine whether particular covariates contributed considerably to the output variable. For model choice, we preferred models with lower WAIC scores. Additionally, we assessed model performance by randomly splitting the data into training and validation datasets and calculating the number of observations correctly predicted by the models. We repeated this process 100 times for each burial side, and for the baseline, full and ‘best’ models, obtaining an average proportion of observations correctly predicted. Results are shown in Figure S3.

We further tested whether culture, rather than genetic ancestry or mobility, can explain the spatiotemporal patterns in burial side. We repeated the Bayesian Latent Gaussian modelling approach implemented in INLA and inlabru on a subset of the data, considering only grave individuals belonging to the 10 most represented cultures, and that also have information on burial side, mobility, and genetic ancestry (n = 755). The data covers a period between ∼8,500 and ∼3,000 calBP. For each burial side (left, right, back), we fit the following sets of models:

1. *Baseline*
2. *Baseline + mobility + ancestry + culture*
3. *Baseline + mobility + ancestry*
4. *Baseline + ancestry + culture*
5. *Baseline + mobility + culture*
6. *Baseline + ancestry*
7. *Baseline + mobility*
8. *Baseline + culture*

Models were parameterised in the same way as described above. Because culture is a categorical variable, INLA and inlabru model it as a random effect. We then focused on the 95% credible intervals of posterior densities of the coefficients for the covariates, and we assessed model performance using WAIC scores. When checking for predictive accuracy of the models, the full model was the one with the worst fit to the data for all burial sides (Figure S4). This is due to potential correlation among explanatory variables and potential overparameterization of the model, making it difficult to interpret the results. We decided to run another set of models where only one variable was added to the baseline, to better investigate the importance of each variable:

1. *Baseline*
2. *Baseline + NEOL*
3. *Baseline + EHG*
4. *Baseline + CHG*
5. *Baseline + WHG*
6. *Baseline + mobility*
7. *Baseline + culture*
8. Variable importance was then assessed by checking the predictive accuracy of the models, based on WAIC scores.

### Modelling the spatiotemporal effects of ancestry, mobility, and culture on burial orientation

We explored whether patterns exist in the orientation of burials and whether such patterns can be best explained by cultural affiliation and/or the genetic ancestry estimates, after controlling for autocorrelation in space and time. Information on burial orientation is available for 1,470 grave individuals. The data consists of a minimum and a maximum angle in degrees from north (0/360°). We calculated the mean between the two using the “mean.circular” function from the “circular” R package (Agostinelli & Lund, 2025). In cases where only the minimum angle was provided, we used that value as the mean. Hereafter we use the mean angle to determine the direction of burial orientation, which refers to the orientation of the head.

We used circular histograms to visualize the data across all individuals. We also grouped the data and visualized patterns across burial sides, period, culture, sex, age, ancestry and mobility. For each variable, we also calculated the pairwise smallest angular distance between groups. Let:

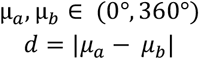

Where μ_*a*_, μ_*b*_ are the circular means within two groups (*a, b*). The smallest angular distance *D*(*a, b*) is:

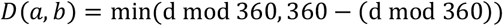

To statistically evaluate differences in burial orientation across cultures, we ran a Bayesian circular regression model, implemented in the “bpnreg” R package (Cremers, 2021). This model can be specified in a way that is mathematically equivalent to a circular ANOVA (Cremers & Klugkist, 2018). Because this is a Bayesian approach, we chose 10,000 output iterations, a burn-in period of 1,000 and we chose to keep a lag every 3 iterations. The model outputs summary statistics for the posterior distributions of the circular means for each culture: the posterior mean, mode, standard deviation (sd) and the lower and upper bound of the 95% highest posterior density interval (HPD) (Cremers & Klugkist, 2018). HPD intervals can be used to test for differences between groups, based on whether the HPDs for the circular mean of the different groups overlap (Cremers & Klugkist, 2018). Because this method tests for differences against a reference group (the Intercept), we computed multiple Bayesian circular regression models while rotating the culture of reference.

This model can also be used with both categorical and linear covariates. To better understand whether space, time, ancestry and culture are good predictors of burial orientation, we tested different regression models with different combinations of covariates:

1. *Spatiotemporal: Burial orientation ∼ Latitude + Longitude + Date*
2. *Spatiotemporal + culture*
3. *Spatiotemporal + ancestry (EHG + CHG + LVN)*
4. *Culture only*
5. *Ancestry only*
6. *Spatiotemporal + culture + ancestry*

In these models, the data used is a subset, as not all grave individuals had information on both burial orientation and ancestry (n = 952). We then looked at the WAIC scores to assess model performance. All analyses were performed using R Statistical Software (v4.6.0; R Core Team, 2026). All data and code used to run the analyses and produce the results are available on GitHub (https://github.com/ecanteri/Burials.git).

## Results

### Burial side and body positioning

We first performed an exploratory analysis of the burial side and positioning values of the grave individuals available in BIAD, for the period and region under study. Inhumation and cremation are the most commonly used deposition types in our data (Figure S1).

Analysis of structure using MCA shows three main groups of body positioning and burial sides. The first cluster consists of the bottom burial side with the scattered, heap, inside a vessel, and sitting body positionings. A second group is represented by the back burial side and the extended body positioning. The left and right burial sides, and their combinations, form the third cluster together with the crouched body position. The front burial side and the prone and irregular body positionings cluster between the back/extended and left/right/crouched groups in MCA space (Figure 3A). This clustering is best understood by looking at the UpSet plot (Figure 3B), which shows that, in our data, individuals buried on their left and right sides tend to also be crouched, while individuals buried on their backs mostly take an extended body position.

This clustering does not seem to show patterns in relation to sex (Figure S7). Instead, we do see patterns in time, where graves from the (early) Bronze Age and Iron Age associate with the sitting/inside a vessel/bottom cluster, together with graves from the Mesolithic and Neolithic, which also associate with the back/extended group (Figure S7). Graves from the Eneolithic, and the Eneolithic/Bronze Age transition pattern with the crouched/left/right cluster (Figure S7). When looking at the ten most represented cultures in our data, we see that individuals from the Lažňany, Linearbandkeramik (LBK), and Baden cultures tend to cluster mostly with the left/right/crouched group, but also with the bottom/inside a vessel/sitting group, while individuals buried in an extended position on their backs belong mostly to Dnieper-Donets, Trichterbecher and Baltic Mesolithic-Neolithic cultures (Figure S7).

The relationship between the identified clusters and genetic patterns remains unclear. Based on the MCA, individuals with higher proportions of WHG and EHG ancestries, and whose estimated spatial distance of maximum genetic similarity (at 250 years earlier) was greater (i.e. high ancestral mobility), tend to have been buried on their front (Figure S8). Higher ancestral mobility is also seen in some individuals within the left/right/crouched cluster and between the left/right/crouched and the sitting/inside a vessel/bottom cluster in MCA space, although the pattern is not so strong (Figure S8). In contrast, a clear pattern is seen with the NEOL ancestry, where individuals with higher proportions of this ancestry belong to the left/right/crouched and the sitting/inside a vessel/bottom cluster (Figure S8). When explicitly modelling the relationship between genetic ancestry, mobility and burial side in space and time, we find that the model with the lowest WAIC score is one in which higher NEOL ancestry contributes to a higher probability that individuals are buried on their left sides, after controlling for spatiotemporal autocorrelation (Figure 4). This is true when modelling across the entire time period (10k – 2k cal BP), and when modelling individuals before and after 5k BP, when Steppe ancestry starts appearing in Western Europe (Figure S9 – S11). When modelling individuals after 5k calBP, we find that the lowest WAIC-scoring model is one in which, additionally, steppe ancestry has a positive contribution to the left burial side (Figure S11).

**Figure 4.**
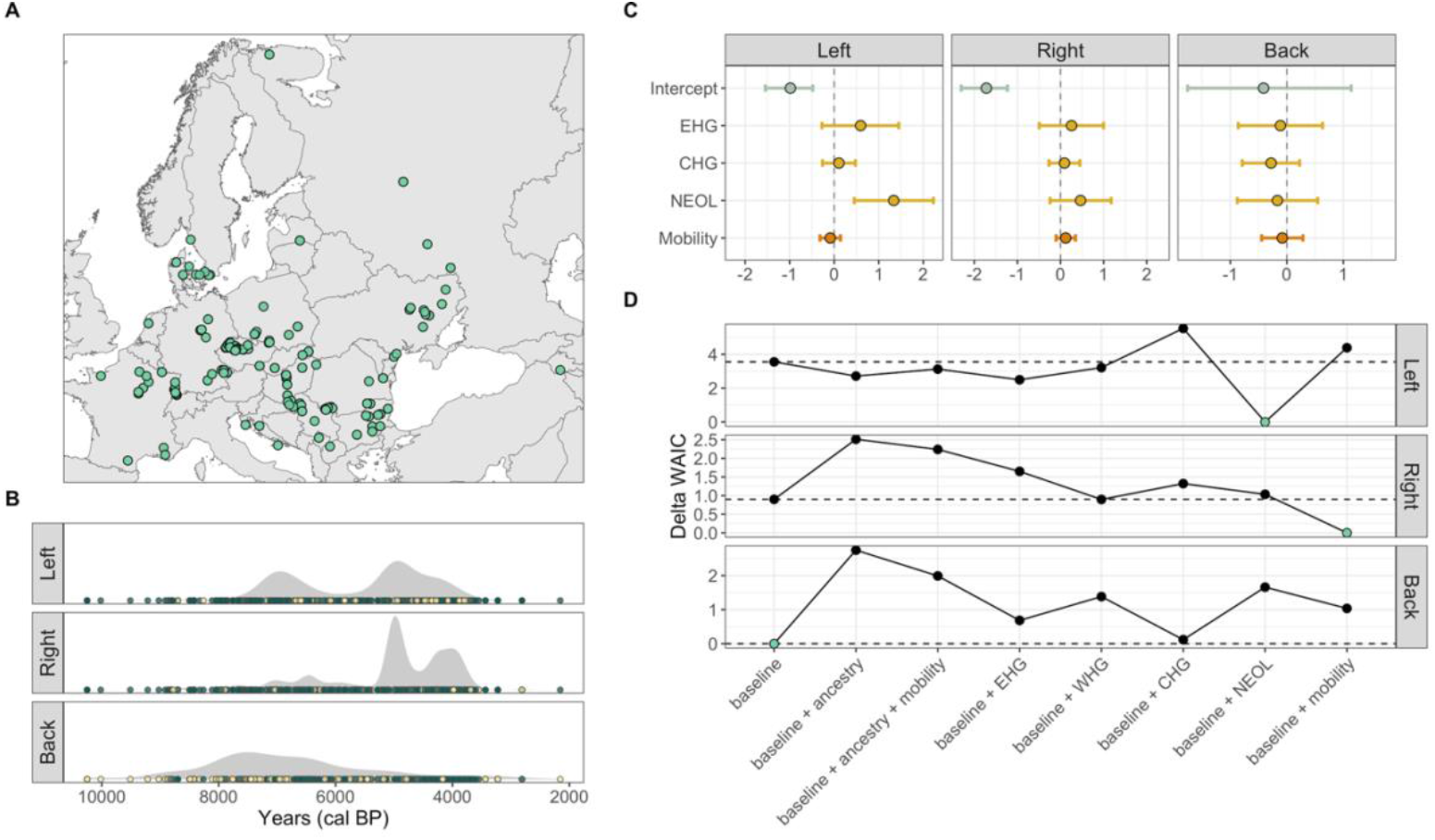
Spatiotemporal relationship of genetic ancestry and mobility with left, right and back burial sides. **A)** Spatial distribution of grave individuals included in the models and **B)** respective radiocarbon age densities for each burial side, with yellow points showing presence (1) of the specific burial side and green points showing absence (0). **C)** Posterior densities of the coefficients for the explanatory variables: the intercept in light blue, Eastern Hunter Gatherer (EHG), Caucasus Hunter Gatherer (CHG), and Neolithic farmers (NEOL) ancestries in yellow, and mobility in orange. The position of the 95% credible intervals is used to identify the effect of the variables on the burial sides **D)** Delta WAIC scores have been calculated by subtracting the minimum WAIC score from all the scores. The dashed horizontal line represents the delta WAIC score for the baseline model (null model). The green points show the models with the lowest score (0), and therefore the ones that have the best fit. Delta DIC and Delta WAIC scores for all models tested, including full models with WHG ancestry, are shown in Figure S12.

Our models do not show an effect of genetic ancestry on the right burial side. When testing multiple models with different combinations of variables, the model with mobility as the only explanatory variable is the one with the best penalized fit to the data (lowest WAIC score), although the the 95% credible intervals for mobility cross zero in the full model (Figure 4). Grave individuals before 5k calBP and buried on their right side show a positive association with mobility, indicating longer geographic distances from their ancestral origin (Figure S9, Figure S10). For the data before 5k calBP, the model that includes mobility is also the one with the best penalized fit, indicating that mobility is a good predictor for the right burial side. This relationship is no longer present after 5k calBP (Figure S9, Figure S11).

The back burial side does not show a relationship with genetic ancestry or mobility, with the baseline model (null model) having the lowest WAIC scores (Figure 4). Only when modelling individuals after 5k calBP, do we see a negative relationship of the back burial side with LVN and steppe ancestry, indicating lower proportions of these two ancestries in grave individuals after 5k calBP buried on their backs (Figure S9, Figure S11).

When adding culture as an explanatory variable, the effect of ancestry on burial side is greatly reduced, at least on the subset of grave individuals we were able to test it on (Figure 5), which is limited in its spatiotemporal distribution. Instead, we see that culture becomes an important variable. Models for the left burial side show that the fit is improved when adding CHG and WHG ancestries, but even more with culture (Figure 5). The null model is the one with the worst fit for the right burial side, and culture is the most important variable. Culture is also the only variable that improves model fit for the back burial side. Still, the effect of each culture varies among burial sides. Dnieper-Donets, Trichterbecher and Baltic Meso-Neolithic cultures show a negative relationship with the left and right burial sides, although their effect is not considerable (the 95% credible intervals of posterior densities cross zero) (Figure 5). All other cultures show a similar pattern for the left and right burial sides, with a positive posterior mean but a 95% credible interval crossing zero (Figure 5). The back burial side is the only one that shows strong effects of cultures, specifically a negative relationship with Lažňany, Únětice and Corded Ware cultures, and a positive relationship with Dnieper-Donets and Trichterbecher cultures (Figure 5).

**Figure 5.**
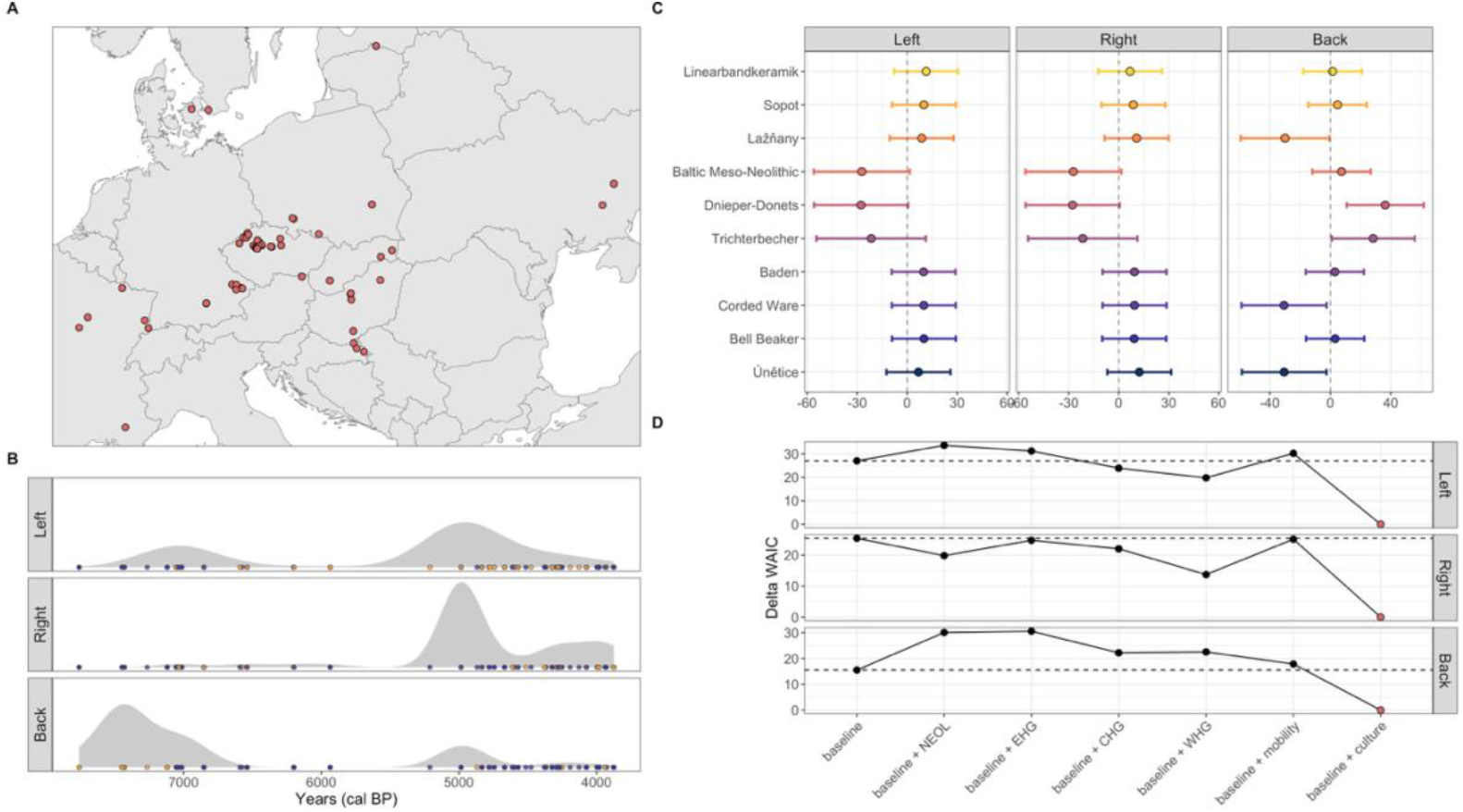
Spatiotemporal relationship of genetic ancestry, mobility, and culture with left, right and back burial sides. **A)** Spatial distribution of grave individuals included in the models and **B)** respective radiocarbon age densities for each burial side, with yellow points showing presence (1) of the specific burial side and blue points showing absence (0). **C)** Posterior densities of the coefficients for cultural affiliation, the random effect in the model with the best fit to the data (baseline + culture). The position of the 95% credible intervals is used to identify the effect of culture on the burial sides. Each colour of the 95% credible intervals represents a specific culture (y-axis). **D)** Delta WAIC scores have been calculated by subtracting the minimum WAIC score from all the scores. The dashed horizontal line represents the delta WAIC score for the baseline model (null model). The red points show the models with the lowest score (0), and therefore the ones that have the best fit. In this set of models, only the spatiotemporal process and one other variable (single ancestries, mobility, culture) have been used as predictors. Results for models using a combination of ancestry, mobility and culture as predictors are shown in Figure S4.

### Burial orientation

The burial orientation of grave individuals covers a wide variety of angles, including all cardinal directions but with a prevalence of east-west (E-W), south-north (S-N) and south-east-north-west (SE-NW) directions (Figure 6). We observe distinct patterns in burial orientation across cultures. Corded Ware and Únětice individuals are buried along a predominantly north (N) - south (S) axis, while Sopot individuals tend to be more aligned along an east (E) – west (W) axis. Baden and Dnieper-Donets individuals prevalently exhibit burials along a SE direction, Trichterbecher and Baltic Mesolithic-Neolithic individuals are predominantly oriented along a south-west (SW) direction, while LBK and Lažňany individuals along an E-SE direction (Figure 6). Differences in cardinal direction are also evident across burial sides, with individuals buried on their left sides oriented mostly E and SE, individuals buried on their right sides oriented SE and S, and individuals buried on their backs oriented NE, E, and S (Figure S14). When studying burial orientations by period, we see that older periods tend to exhibit a S orientation, while later periods show a shift towards N and NE directions (Figure S15). We do not find pronounced differences in orientation between male and female burials (Figure S16).

**Figure 6.**
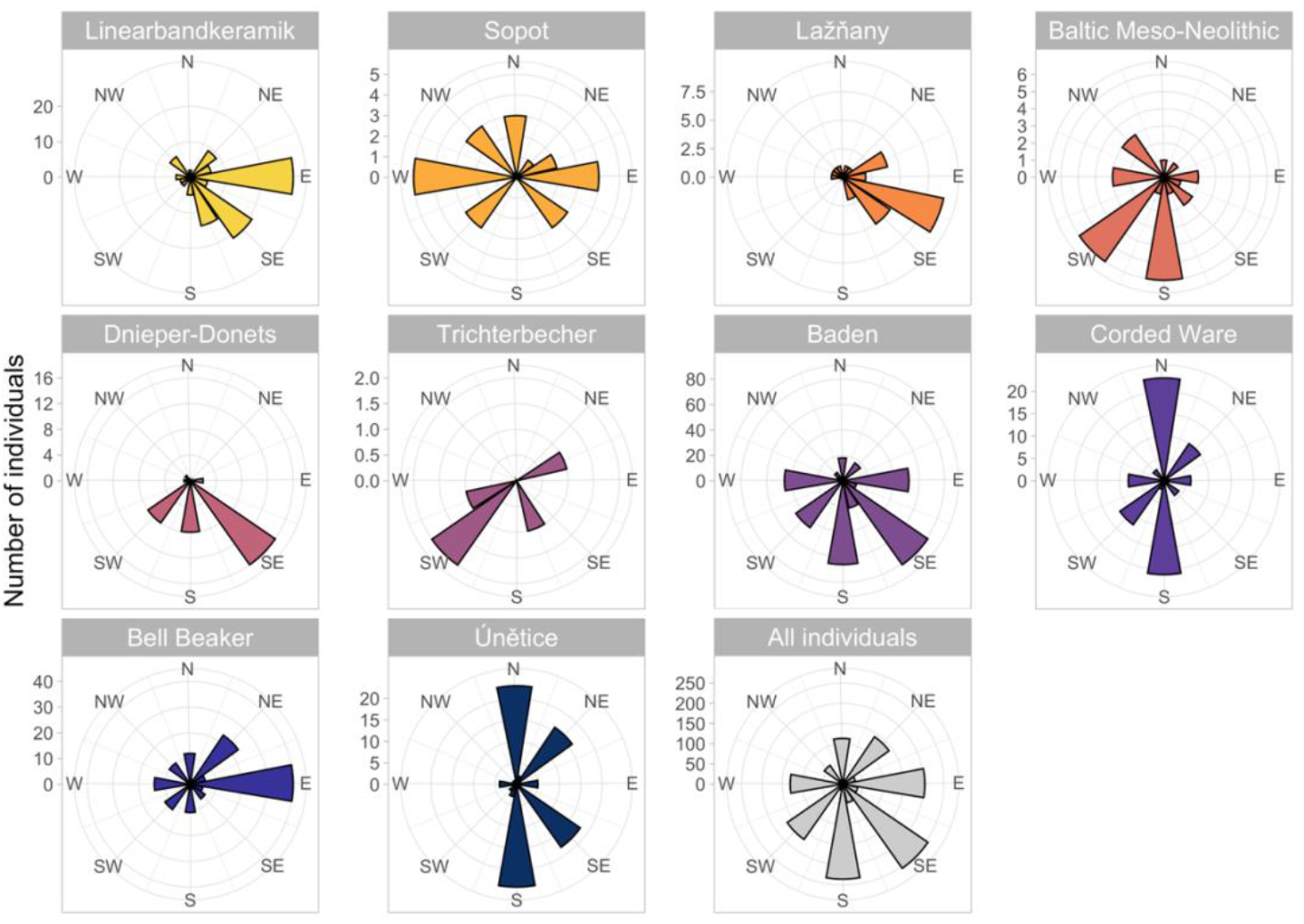
Burial orientation across cultures. Histograms show the number of individuals and their burial orientations, for graves belonging to the ten most represented cultures in our data, from oldest to youngest. Shown here are only individuals for which we have both cultural and ancestry information. The orientations of all individuals belonging to the same cultures are shown in Figure S13.

The Bayesian circular regression analysis (Figure 7) allows us to test for differences in circular mean across cultures. This method estimates the 95% highest posterior density interval of the orientation distribution for each culture and then specifically tests for pairwise differences among the means. If the upper and lower bounds of the posterior distributions cross zero, then the two means are considered “not statistically different”, while if the converse is the case, the means are considered “statistically different”. Based on these cutoffs, Bell Beaker individuals are statistically different from LBK and Lažňany individuals, and LBK, Lažňany, Bell Beaker, and Únětice individuals have statistically different circular means compared to Baden, Dnieper Donets, Trichterbecher and Baltic Mesolithic-Neolithic individuals (Figure 7, Table S1). Among Baden, Dnieper Donets, Trichterbecher and Baltic Mesolithic-Neolithic cultures, only Baden and Baltic Mesolithic-Neolithic individuals have statistically different circular means (Figure 7, Table S1). Corded Ware individuals are only statistically different from individuals belonging to Bell Beaker and Lažňany cultures, and statistical difference in burial orientation also results between Sopot, Baden, and Dnieper-Donets cultures (Figure 7, Table S1).

**Figure 7.**
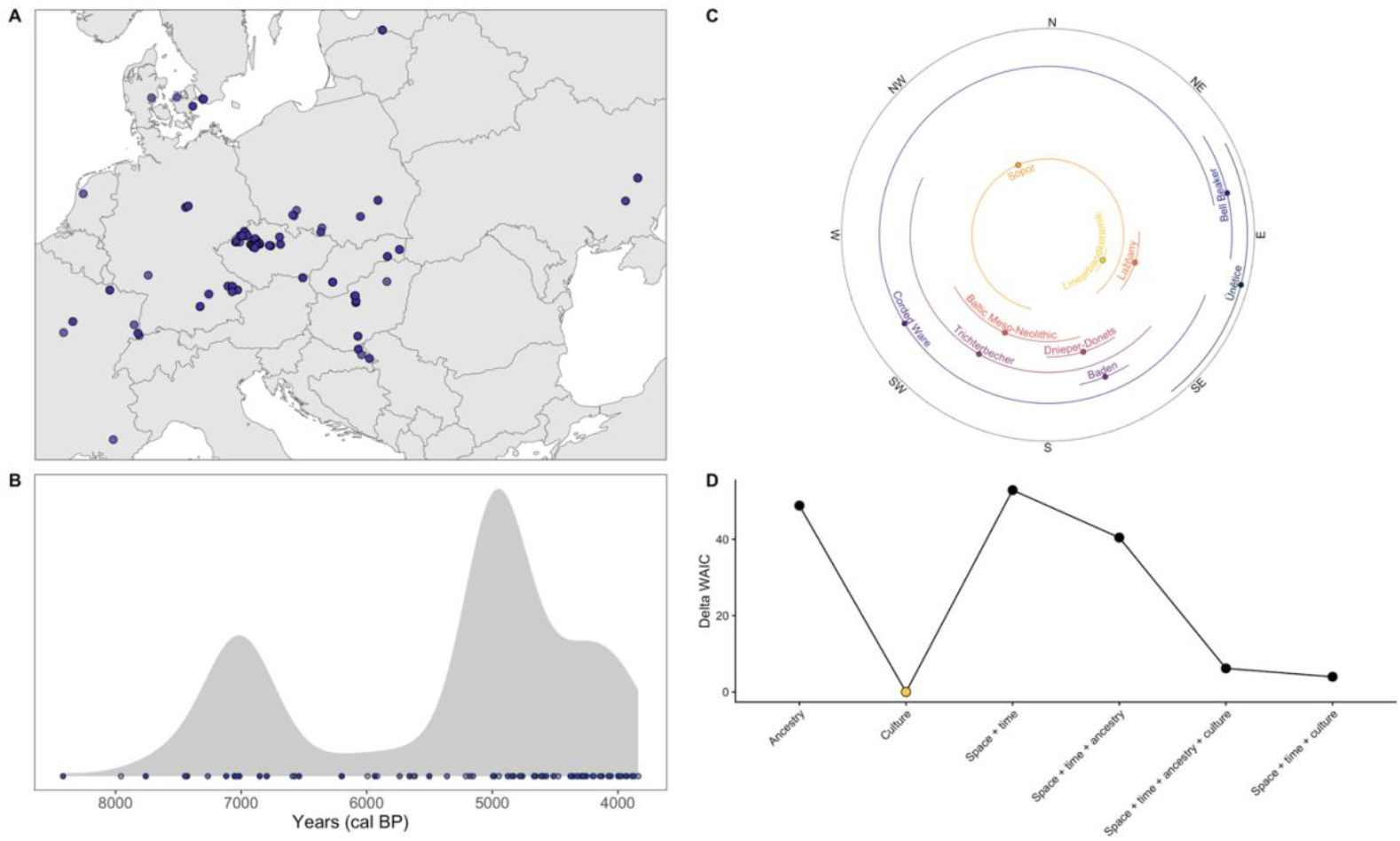
Differences in burial orientation across cultures. **A)** Spatial distribution of grave individuals belonging to the ten most represented cultures in our data, and with information on burial orientation and genetic ancestry. **B)** Their respective radiocarbon age density. **C)** Results of the Bayesian circular regression testing for differences in burial orientation across the ten most represented cultures in our data. Shown are the posterior distributions of the circular means for each culture, with the points representing the mean of the distribution and the lines the upper and lower bounds. Cultures have statistically different circular means if their posterior distributions do not overlap. Only individuals with both ancestry and culture information were included in the models. Results of the Bayesian circular regression including all individuals are shown in Figure S17. **D)** Delta WAIC scores for the tested models. Delta WAIC has been calculated by subtracting the minimum WAIC score from all the scores. The gold point shows the model with the lowest score (0).

After testing multiple models with different combinations of explanatory variables (space, time, ancestry, and culture), the models with the lowest WAIC scores were by far the ones that included culture (Figure 7), indicating that culture is an important variable for explaining burial orientation.

## Discussion

In this paper we have attempted to approach debates concerning the relationship between archaeological cultures and clusters of genomic variation as revealed by aDNA studies by using explicitly spatial approaches, as well as dimensionality reduction methods. As palaeogenomics routinely relies on archaeological cultures to contextualize the findings (for a different approach cf. Fu et al., 2016), one of our goals was to determine whether genetic affinity can account for cultural attributes, meaning that genetic and cultural transmission run in parallel. To this end we modelled different factors that could have affected patterns in two attributes of ancient inhumation burials: the body position of the buried individual and the orientation of the burial with respect to the points of the compass. Although both are routinely published as cultural attributes, they are often treated as secondary for assigning cultural classification and their values do not have inbuilt associations with specific cultures.

Burial rituals are generally regarded as conservative traits subject to the social norms of specific communities (Binford, 1971; Saxe, 1970; Ucko, 1969). Potentially, this might mean they could be influenced by patterns of vertical inheritance if rituals are handed down from parents to children or within a given community to descendant communities, which could, in turn, potentially lead to associations with genetic ancestry (Mesoudi, 2011; Richerson & Boyd, 2005). Burial rituals can also be influenced by social norms or beliefs in other domains that may or may not run parallel with genetic ancestry. Randsborg and Nybo (1984) suggested a re-occurring temporal pattern between body orientation and sunset position on the day of burial in Early Bronze and Viking Age burials. Carr (1995) indicated that features connected with burial positioning seem to be influenced by beliefs about the universal order and the afterlife. A quantitative study of burial orientation in Japanese jar burials (Maikuma & Nakao, 2024) addressed the hypothesis that the northerly direction was influenced after the Kofun (early historic) period by Confucian or Buddhist ideas. They proposed that this would be more plausible if burials in the previous period were not oriented in this direction and found this to be the case. A recent study of the relationship between language affiliation and patterns of variation in mortuary practices among groups speaking Pama-Nyungan languages in Australia (Learmouth et al., 2024) found some association between the two. However, this was not nearly as strong as that for initiation rituals.

In this study, we modelled the effect of genetic ancestry, genetic mobility and cultural affiliation on the selected burial attributes, while controlling for spatiotemporal autocorrelation. Our results have shown a range of complex and varied patterns, but in virtually all cases there was evidence of population structure going beyond isolation-by-distance in space and time. Similarly, when Schmid (2020) compared time trends of cultural similarity between burials, in terms of whether they were inhumations, cremations, flat graves, or graves with mounds, with their spatial distance, he found that there was ‘no significant permanent correlation’, suggesting that their spread could not be accounted for by simple diffusion.

For the models that did not include culture, higher NEOL ancestry, associated with the spread of the first farmers into Europe, contributes to a higher probability that individuals were buried on their left sides. However, after 5k calBP, we find that the best model is one in which, additionally, steppe ancestry, derived from the westward spread of groups with ancestry in the Eurasian steppe, has a positive contribution. Perhaps surprisingly, we do not see a corresponding effect of genetic ancestry on the right burial side, where mobility is the only factor that matters. As explained above, mobility refers to the distance between the place where the individual was buried and the estimated spatial location of the point of maximum genetic similarity to that individual 250 years earlier (Schmid & Schiffels, 2023). The implication of the result is that those individuals buried on their right-hand side, or their immediate ancestors, may have a history of greater directional mobility, but at this point it is not clear why this should be the case. For supine (back) burial placement, the baseline space-time distance model is the best fit. However, for the more limited dataset where cultural affiliation information was available in addition to ancestry and mobility values, its addition as an explanatory variable resulted in considerable reduction in ancestry effects. Now the left- and right-side burials are best modelled with ‘culture only’ and the same is true of burial on the back. Nevertheless, for the great majority of cultures, the 95% credible intervals of the coefficients of the best models include zero. Supine burial is the only one that shows a strong effect of cultures, negative in the case of Lažňany, Únětice and Corded Ware cultures, and positive for the Dnieper-Donets and Trichterbecher cultures (Figure 5).

In the case of burial orientation, the results are more clear-cut. After comparing multiple models with different combinations of space, time, ancestry, and culture as explanatory factors, the best models by far were ones that included the archaeological culture to which the burial was assigned by the archaeologists writing the report. This indicates that archaeological culture is an important variable for explaining variation in burial orientation. In fact, the very best model only included culture; in other words, if we know the culture to which a burial is assigned, we can make a better prediction of its orientation than using ancestry, geographic or chronological data. Thus, the present case-study confirmed the long-standing observation that the orientations of Corded Ware burials were significantly different from Bell Beaker ones, with which they overlap in space and time, despite data aggregation irrespective of the biological sex and strongly gendered burial rites (Bourgeois & Kroon, 2017; Strahm, 2002; Wentink, 2020). In comparison to Corded Ware, the narrow distribution of the orientation range for Bell Beaker burials suggests the importance of high-fidelity transmission of burial norms as well as the signalling of difference. These two cultures are closely related in terms of their genomic ancestry, but with distinct Y-chromosome signatures. We note, however, that genetic factors in our study are summarized into coarse ancestry scores, and we cannot discard the possibility of influence of more fine-scale genetic factors (which may not be liable to be captured in a simple SNP-based ADMIXTURE analysis) on burial orientation.

## Conclusions

We set out to investigate the factors affecting spatiotemporal population structure in an archaeological dataset of inhumation burials in the light of recent heated but inconclusive debates about the relation between archaeological cultural patterns and recent evidence for shifting populations provided by aDNA. Our approach was based on breaking down culture into specific elements related to different domains, following previous ethnographic cultural evolution studies. In our case this involved aspects of burial practices that it is reasonable to believe were governed by social norms, and then modelling factors that could account for them, including genomic ancestry and cultural affiliation, after accounting for spatiotemporal autocorrelation, a significant advance on previous approaches to such questions. Our results showed that SNP-based genomic ancestry scores improved model fit in some cases but by no means all, and that archaeological culture is important, separately from genomic ancestry. This suggests that archaeological cultures can be considered entities, not just archaeologists’ fictions, and that they are not reducible to coarse measures of genomic ancestry. However, they should not be considered as well-defined aggregations of artefacts and contexts linked to a people, i.e. the ‘traditional perception’. Rather they are a kind of summary of the effects of parallel and overlapping transmission networks in different domains that have some separation in space and/or time, as per Clarke’s (1968) polythetic definition of archaeological cultures. What is necessary in the future is to explain how and why different traits are linked in transmission more closely than others, a task which requires more than establishing that the networks summarised as an ‘archaeological culture’ are distinct from those arising through genetic transmission.

However, the genetic clusters considered here are taken from estimates of SNP-based ancestry inference (the WHG, EHG, CHG and NEOL components). Such clusters are rather broad and will miss more local patterns of genetic relatedness, and so a further avenue of analysis would involve using more fine-scaled measures of genomic affinities (including statistics based on sharing of large identity-by-descent segments or tree sequence inference). Additionally, to extend this approach to other archaeological domains, it will be necessary to develop culturally agnostic descriptions of artefact attribute variation within a population variation framework that moves on from thinking in terms of essentialist archaeological types, and whose chronology is based on independent dating. Overall, this study has shown that using explicitly spatiotemporal statistical models that combine information from both archaeological and paleogenetic studies can shine a light on the potential driving factors behind patterns of burial practice variation.

## Acknowledgements

We are grateful to Kristian Kristensen for comments on an earlier draft though we have not always followed his suggestions. This research is part of the synergy project ‘COREX: From Correlations to Explanations: towards a new European prehistory’, funded by the European Research Council (ERC) under the European Union’s Horizon 2020 research and innovation programme (Grant Agreement No. 951385). M.G.T. is supported by ERC Horizon 2020 research and innovation programme grant agreements: no. 951385 (COREX) awarded to M.G.T., no. 865515 (SUSTAIN) awarded to M.I-B., no. 324202 (NeoMilk) awarded to Richard Evershed, no. 788616 (YMPACT) awarded to Volker Heyd, by Wellcome Senior Research Fellowship Grant 100719/Z/12/Z awarded to M.G.T., and by UK Natural Environment Research Council ‘Pushing the frontiers of environmental research’ grant NE/X01469X/1 awarded to M.G.T. M.I-B. has received funding from the European Research Council (ERC) under the European Union’s Horizon 2020 research and innovation programme (Grant agreement No. 865515 SUSTAIN). R.S. was additionally supported by the NAWA Polish Returns 2023 project (BPN/PPO/2023/1/00013/U/00001) and the Polish National Science Center Research Component (2024/03/1/HS3/00008).

