## Supplementary Information for "Cultural affiliation accounts for most of the spatiotemporal variation in burial rite practices"

**Supplementary Methods**

*Body position*

The two traits refer to the archaeologically described arrangement of the discovered deceased. Combined, both are considered culturally significant as they represent an enactment of culture-specific norms of post-mortem transformation of the decomposing tissues (Parker Pearson, 1999, p. 54; Gramsch, 2013, pp. 462–463). Historically, European archaeologists have first used body position and burial side as supplementary variables to material culture in order to identify archaeological cultures. Later, both traits were used to apply the archaeological culture classification to burials without finds (Häusler, 2001). In some instances like the Corded Ware or Bell Beaker, the strength of this relationship has been continuously re-affirmed throughout the last 100 years (Strahm, 2002, p. 184; Furholt, 2019) even if some degree of variability is recognized (Kolář, 2018, pp. 70–72).

In instances where the burial rite, preservation conditions, and the involvement of an osteoarchaeologist allow in-depth characterisation of the burial position, specific information about the body manipulation and deterioration conditions can be documented (Tõrv, 2016; Müller-Scheeßel and Hukel’ová, 2020; Knüsel and Schotsmans, 2022). However, for the vast majority of Western Eurasian burials discovered over the past 100 years, this information is often unavailable unless the preserved remains are subject to a re-analysis accompanied by a detailed examination of the excavation reports (e.g. Fokkens *et al.*, 2017). Where these conditions were not met, the resulting report often focuses on reporting the obvious traits, subject least to any form of post-depositional displacement. This often means a brief statement about the overall position of the deceased, e.g. crouched, sitting or extended with additional details about the rotation of the head of the deceased or the arrangement of the legs (Frînculeasa, Preda and Heyd, 2015).

The heterogeneity of reporting standards required decision-making at the time of data collection to generate a robust dataset with wide spatio-temporal coverage. We decided to record information on body positioning in two independent columns: *BodyPositioning* and *BurialSide*. The former stores the overall position of the body according to the common archaeological standards (crouched, inside a vessel, scattered, extended, head, prone, irregular, sitting, semi-crouched), while the latter reports the categorical rotation axis for the individual. In instances where this information was provided in the examined source, the original classification was entered into the databases. In cases where this information was not available in the text but was available on figures, the information was extracted by our researchers. In cases where different linguistic conventions were used, they were translated into our default English vocabulary and entered into the database.

*Burial orientation*

Burial orientation refers to the direction the deceased is placed towards. This cultural trait is associated with cultural norms dictating that the body of the deceased individual should be directed towards important physical or cosmological landmarks (Ucko, 1969; Saxe, 1970; Binford, 1971). Routinely recorded in archaeological practice and supported by ethnographic records, this trait has received less scholarly attention than body positioning analysis, as accurate identification of the said landmark or its cosmological rationale remains enigmatic. However, from a more contemporary perspective, the prevalence of body orientation norms in two Abrahamic religions - Christianity and Islam - underscore the relevance of recognizing burial orientation as an important cultural variable (Ucko, 1969; Bullion *et al.*, 2022). Similarly to the body position or burial sides, instances where the patterning of orientation rules specific to cultural groups or genders has been well-documented in Europe (Kolář, 2018, p. 72), there are ample cases where no specific rule has been observed (Czekaj-Zastawny, 2021; Müller-Scheeßel *et al.*, 2021).

Two standards of reporting body orientation can be encountered in the literature: cardinal directions-based and angular. The former reports the axis the body was placed alongside, where one of the directions, usually the first, represents the direction the head is pointing towards, while the feet correspond to the other (Müller-Scheeßel and Hukel’ová, 2020; Czekaj-Zastawny, 2021). The latter uses the same logic but rather than providing a rough estimate of the first direction, an azimuth from the north is reported as a numeric value 0-360˚ (Chen, 2025).

For the purpose of our research we decided to use only the second system and in order to incorporate data from case studies, where only the cardinal directions were reported, a conservative confidence interval was used to provide an estimate of the azimuth. In cases where the deceased was reported alongside an image representing the individual *in situ* the image was extracted from the pdf, projected into an *Inkscape* drawn compass rose with 10˚ resolution and the minimum and maximum values were recorded to account for variation in the orientation (Klein *et al.*, 2025). For cases where no visual cues could be obtained a 90˚ confidence interval was for the reported first angle to account for possible errors, measurement imprecision, and general uncertainty (Klein *et al.*, 2025). Where some degree of precision for the sample could be obtained the confidence interval would be reduced after examining the accuracy of the conversion.

**Supplementary Figures**

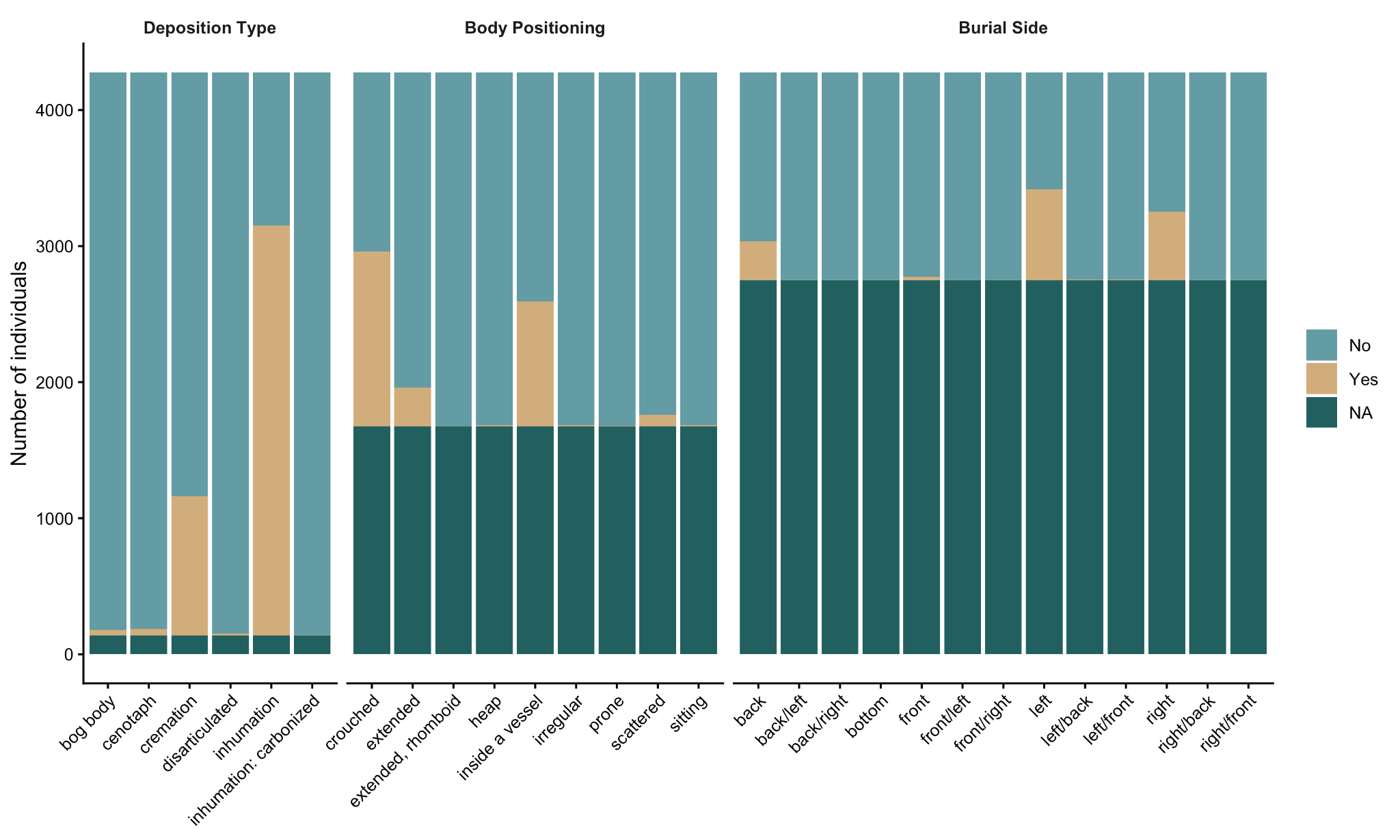

**Figure S1.** Number of grave individuals in our dataset with information about the type of deposition, the position of the body in the burial, and the side on which the individual was buried. Each bar represents a variable in each trait, and the colours indicate the number of individuals associated (Yes), not associated (No) to that variable, or with no information (NA).

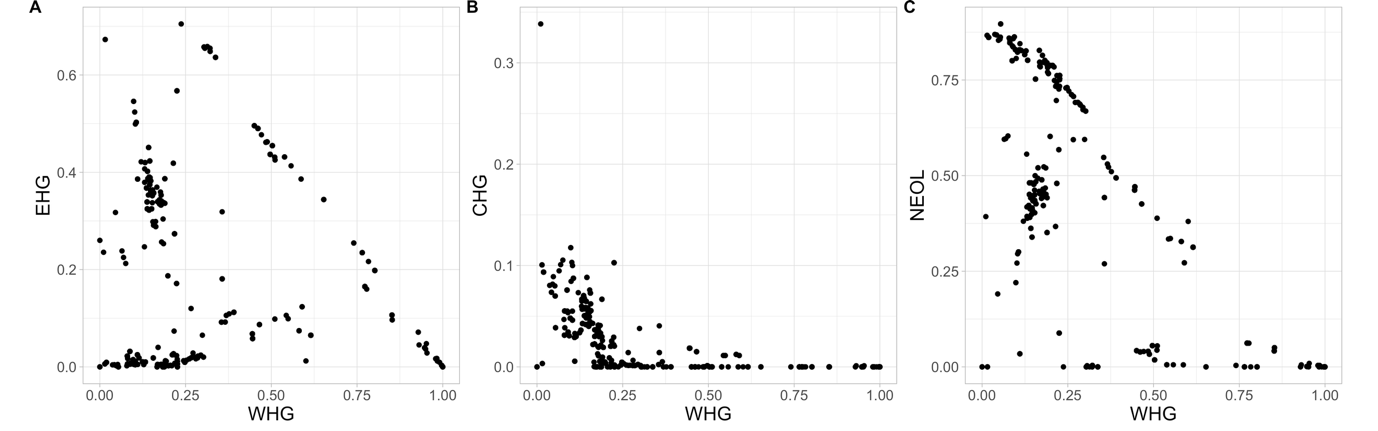

**Figure S2.** Collinearity among ancestry values and the other ancestries in our data. Ancestry values correspond to a broad-scale clustering analysis, and we specifically focused on four ancestries that are maximized among, respectively, Neolithic farmers migrating from Anatolia (NEOL), Western hunter-gatherers (WHG), Eastern hunter-gatherers (EHG) and Caucasus hunter-gatherers (CHG). We specifically tested for collinearity between WHG ancestry (x-axis) and the other ancestries (y-axis): **A)** EHG, **B)** CHG and **C)** NEOL.

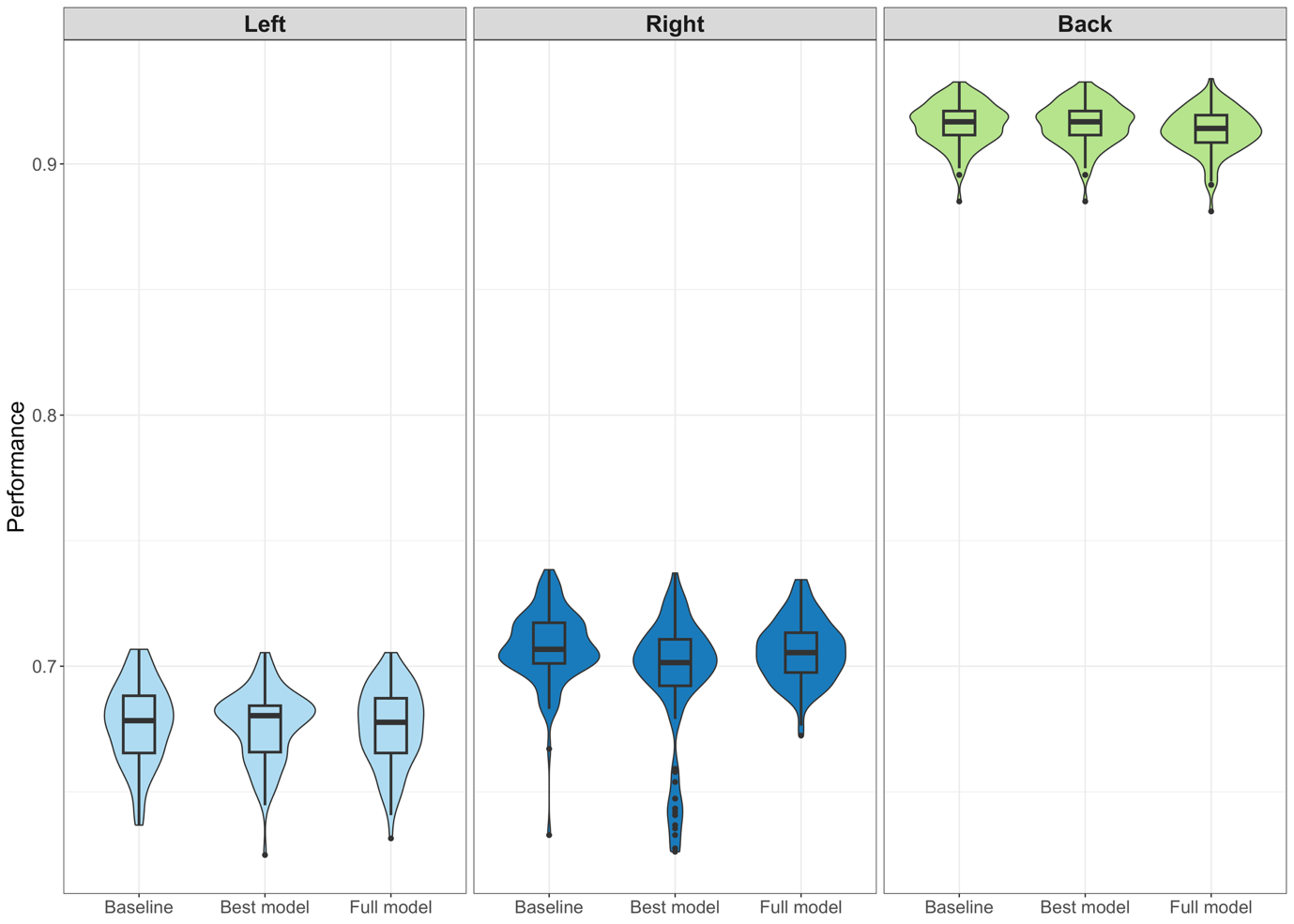

**Figure S3.** Performance of the INLA spatiotemporal models for burial side. We assessed model performance by randomly splitting the data into training and validation datasets and calculating the number of observations correctly predicted by the models. We repeated this process 100 times for each burial side, and for the baseline model, the model with all predictors (full model) and the model with the lowest WAIC score (best model), obtaining an average proportion of observations correctly predicted.

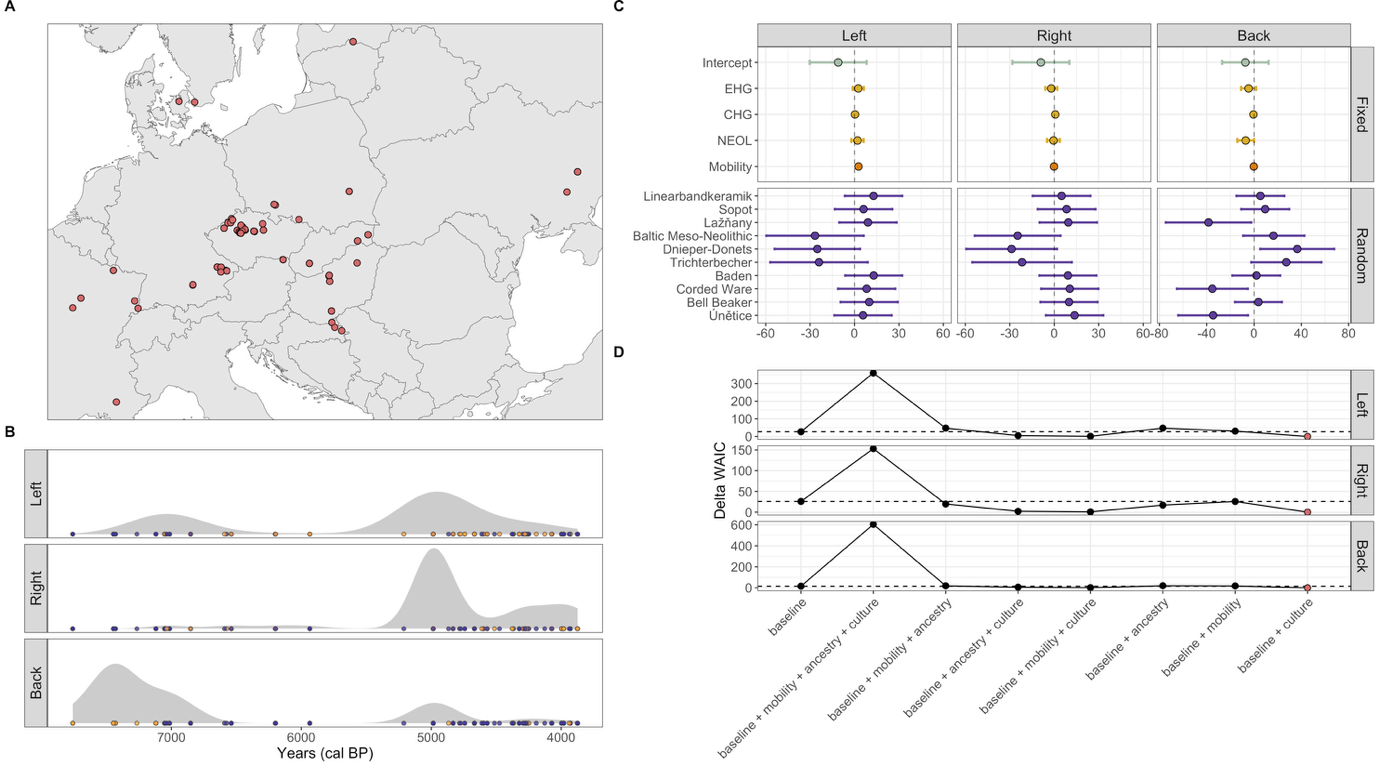

**Figure S4.** Spatiotemporal relationship of genetic ancestry, mobility, and culture with left, right and back burial sides. **A)** Spatial distribution of grave individuals included in the models and **B)** respective date densities for each burial side, with yellow points showing presence (1) of the specific burial side and blue points showing absence (0). **C)** Posterior densities of the coefficients for cultural affiliation, the random effect in the model with the best fit to the data (baseline + culture). The position of the 95% credible intervals is used to identify the effect of culture on the burial sides. Each colour of the 95% credible intervals represents a specific culture (y-axis). **D)** Delta WAIC scores have been calculated by subtracting the minimum WAIC score from all the scores. The dashed horizontal line represents the delta WAIC score for the baseline model (null model). The red points show the models with the lowest score (0), and therefore the ones that have the best fit.

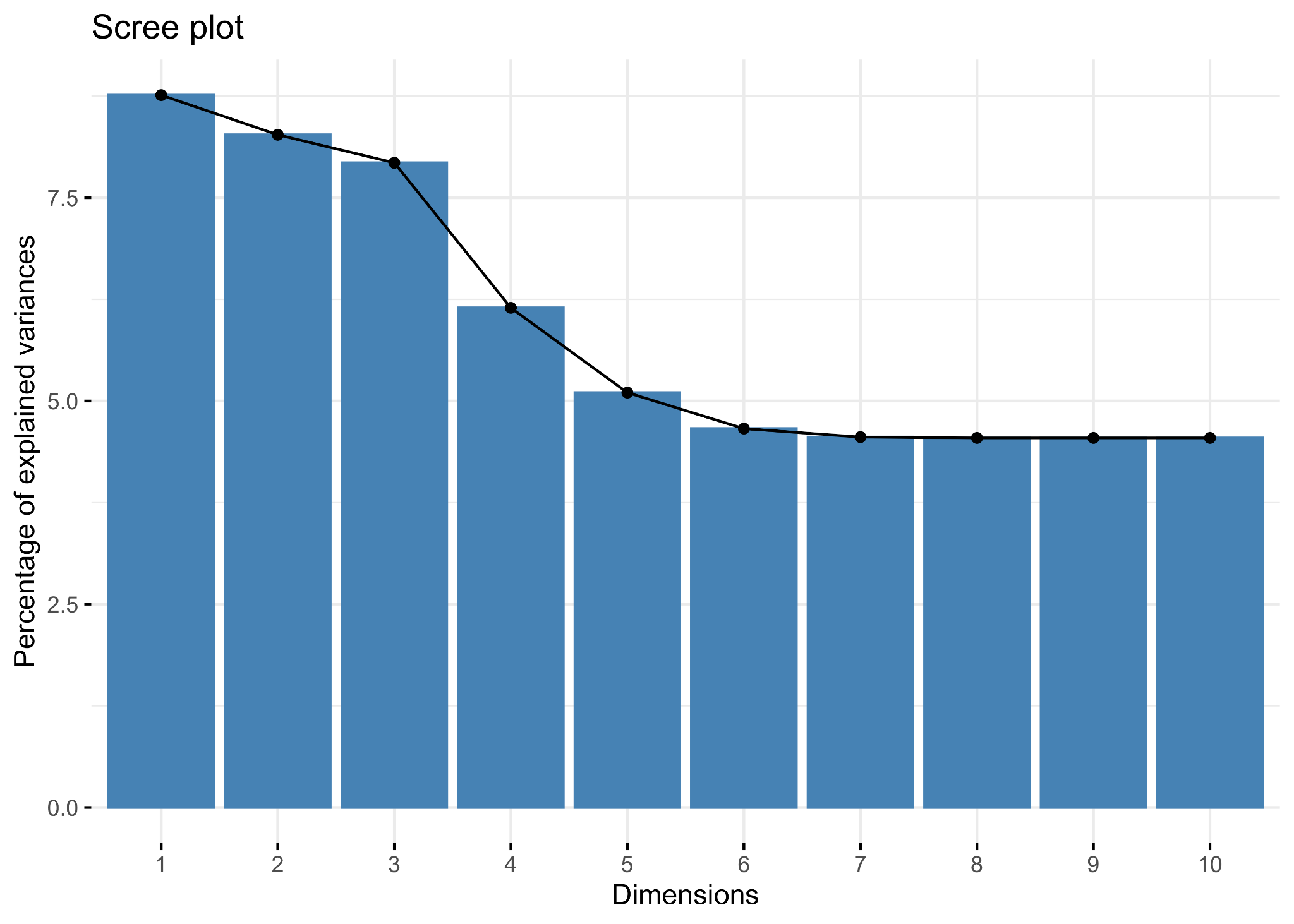

**Figure S5.** Scree plot showing the percentage of variance explained by the 10 dimensions in the Multiple Correspondence Analysis (MCA), for burial side and body positioning of the grave individuals in our data.

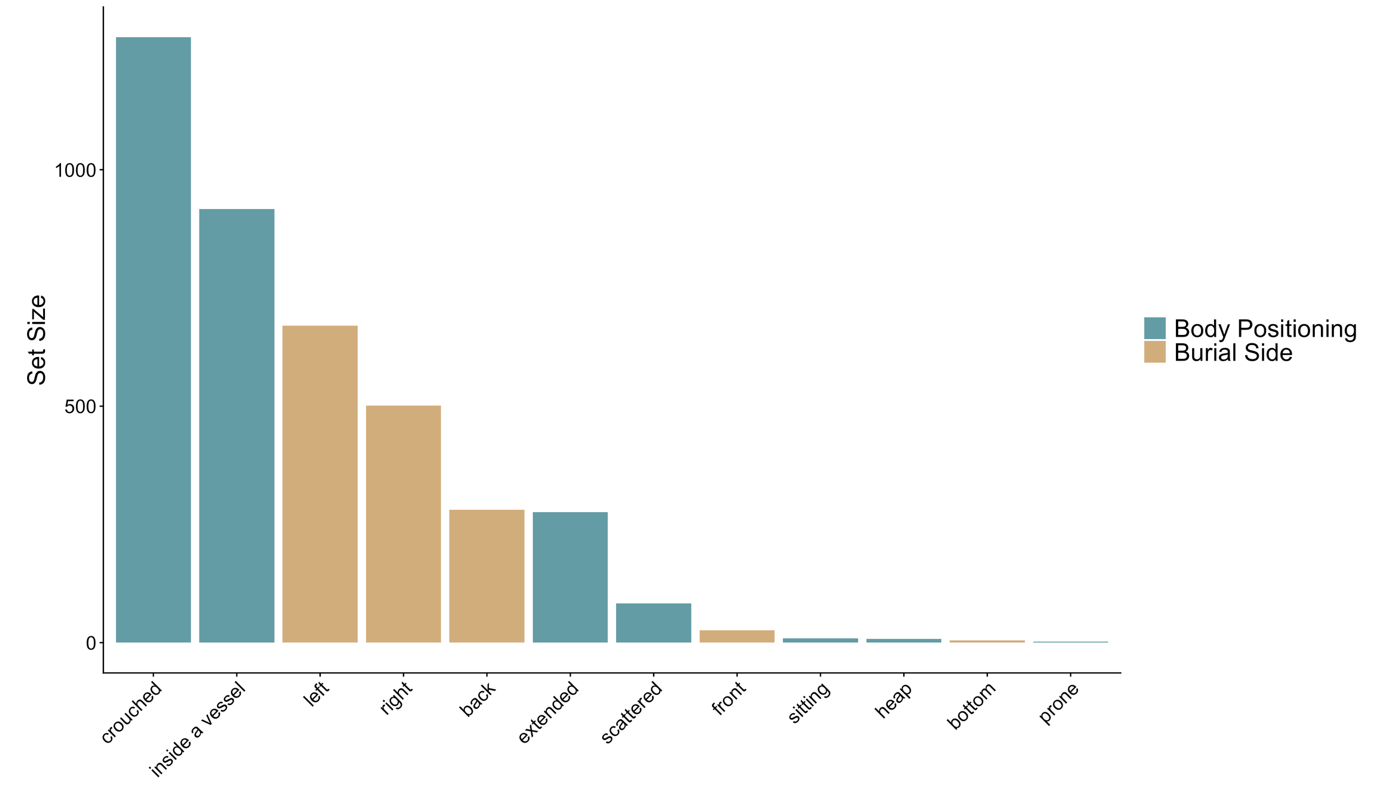

**Figure S6.** Set size in burial side and body positioning. Bars show the number of individuals within each variable (set size) for body positioning (blue) and burial side (yellow).

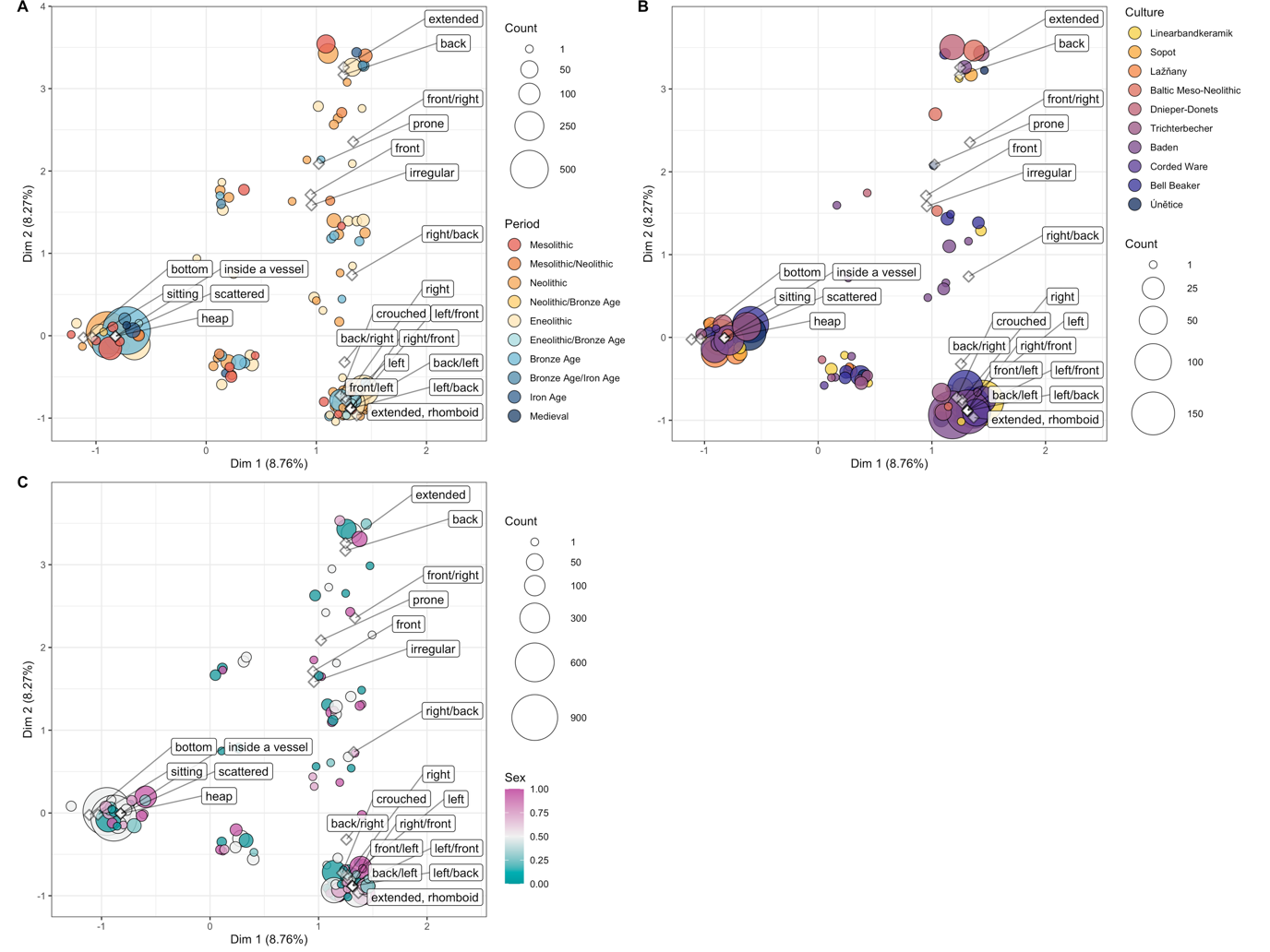

**Figure S7.** Structure in burial rite traditions across period, culture and sex. Multiple Correspondence Analysis (MCA) showing clustering in burial side and body positioning of grave individuals, based on archaeological period (**A**), culture (**B**) and sex of the buried individual (**C**). The size of the circles represents the number of individuals having the same coordinates in the MCA space. Only individuals belonging to the ten most represented cultures in our data have been included in the analysis. The sex of the individual represents the probability of being female based purely on the anthropoplogical (or biological) evidence, without considering material culture. A value of 1 indicates female with certainty, a value of 0 indicates male with certainty, and a value of 0.5 indicates an equal probability of male or female. In cases where the individual has not been osteometrically assessed, we assumed a prior of 0.5.

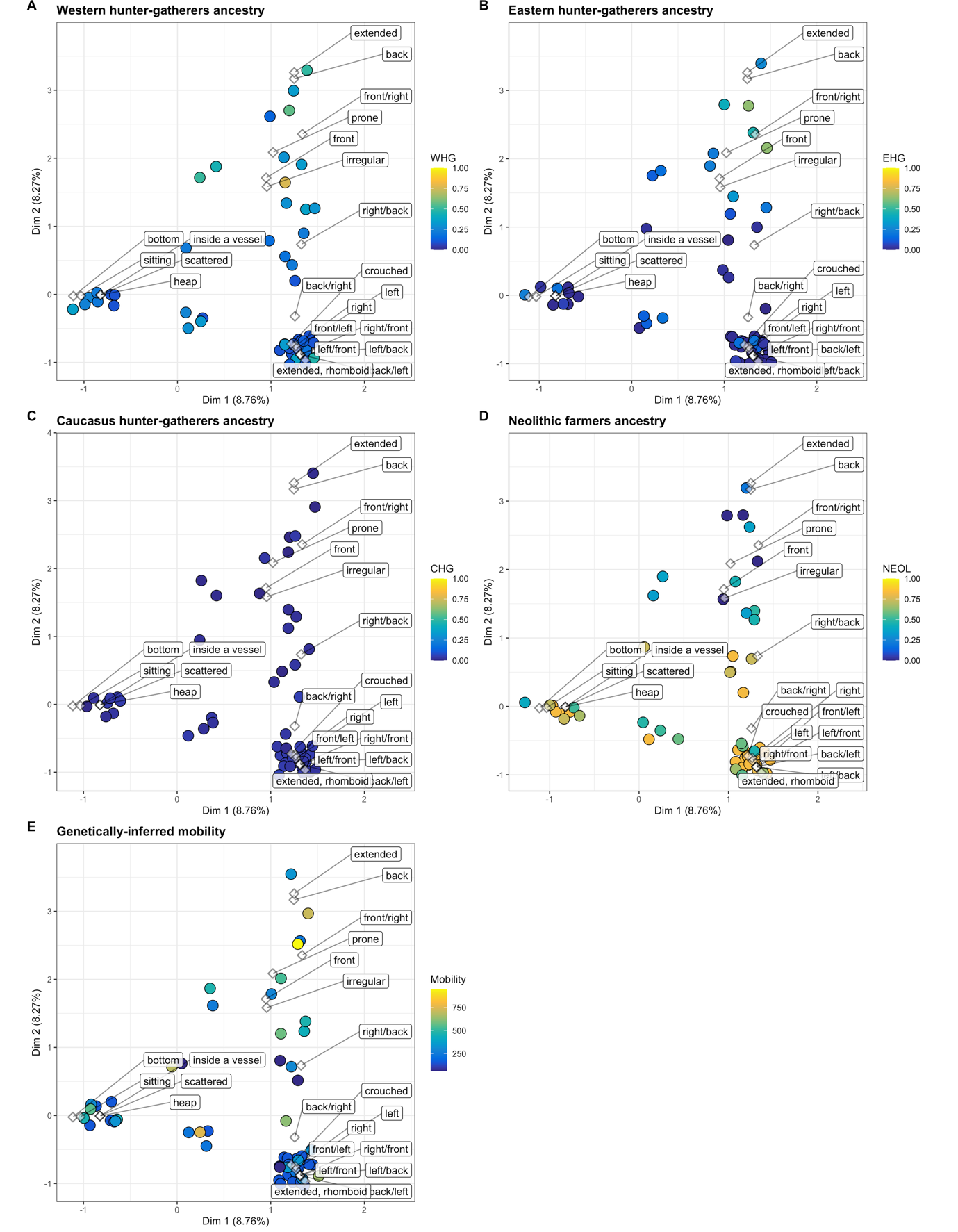

**Figure S8.** Structure in burial rite traditions across ancestry groups. Multiple Correspondence Analysis (MCA) showing clustering in burial side and body positioning of grave individuals, coloured by the genetic ancestry and mobility values associated to them: **A)** Western hunter-gatherer anccestry (WHG); **B)** Eastern hunter-gatherer ancestry (EHG); **C)** Caucasus hunter-gatherer ancestry (CHG); **D)** Neolithic farmers ancestry (NEOL); **E)** genetically-inferred mobility. The size of the circles represents the number of individuals having the same coordinates in the MCA space.

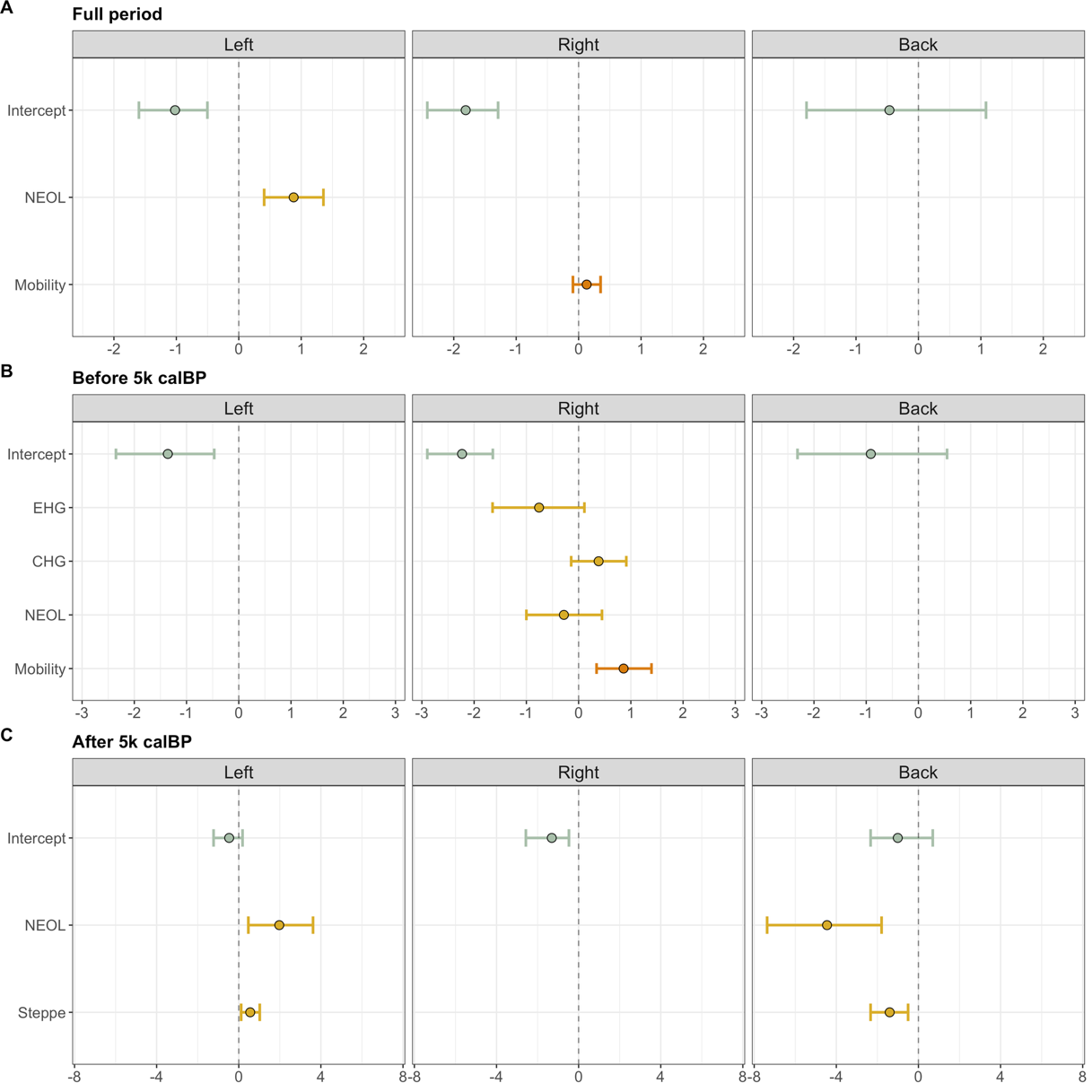

**Figure S9.** Spatiotemporal relationship of genetic ancestry and mobility with burial side across different time periods, resulting from the models with the lowest WAIC scores. Posterior densities of the coefficients for the explanatory variables: the intercept in light blue, Eastern Hunter Gatherer (EHG), Caucasus Hunter Gatherer (CHG), Neolithic farmers (NEOL), and Steppe (EHG + CHG) ancestries in yellow, and mobility in orange. The position of the 95% credible intervals is used to identify the effect of the variables on the left, right and back burial sides. **A)** Posterior densities of the coefficients used in the models with the lowest WAIC scores analysing the full time period (10k – 2k calBP). **B)** Posterior densities of the coefficients used in the models before 5k calBP, for the models with the lowest WAIC scores. **C)** Posterior densities of the coefficients used in the models after 5k calBP (when the Steppe ancestry starts appearing in Western Europe), for the models with the lowest WAIC scores.

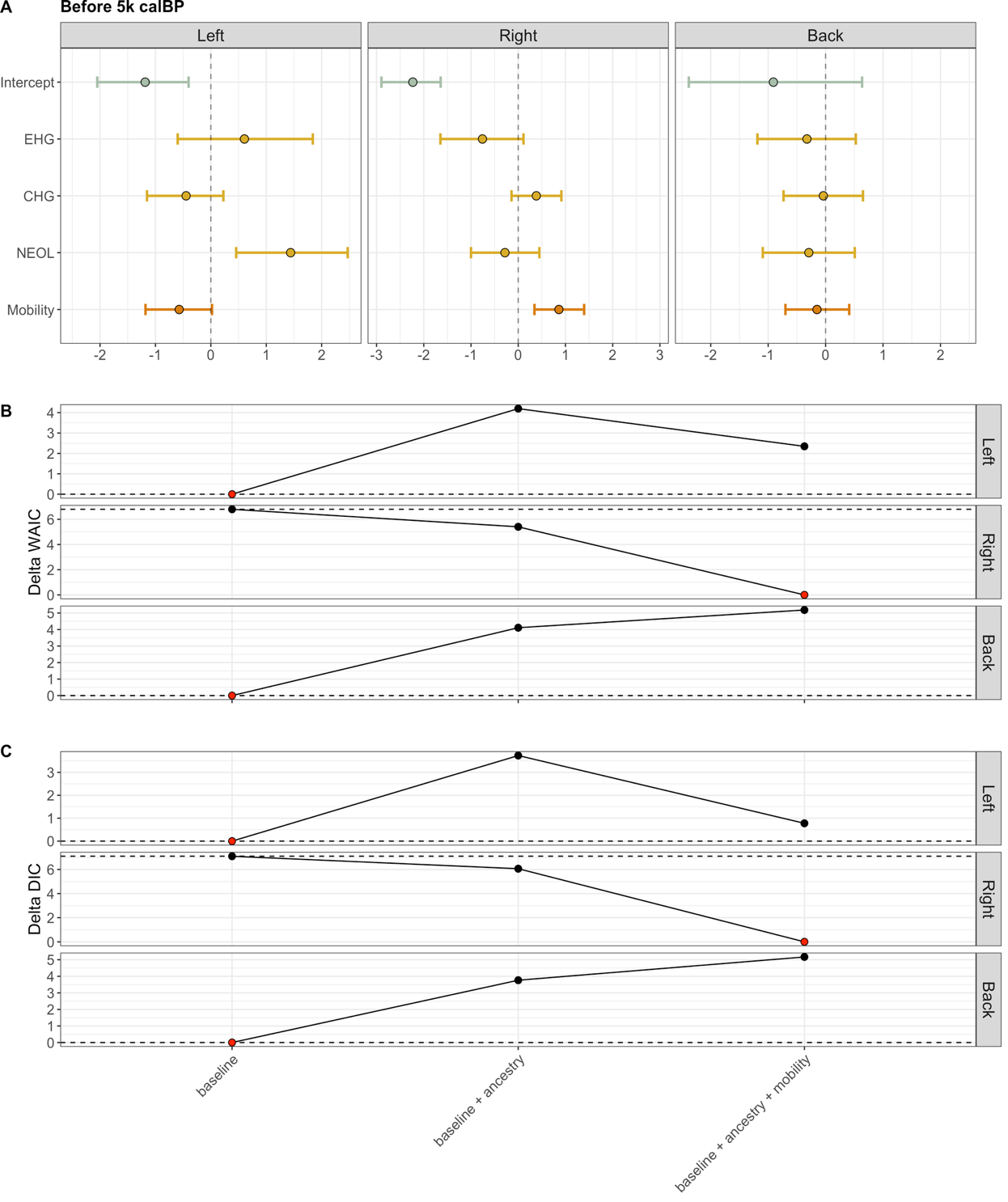

**Figure S10.** Spatiotemporal relationship of genetic ancestry and mobility with burial side before 5k calBP. **A)** Posterior densities of the coefficients for the explanatory variables: the intercept in light blue, Eastern Hunter Gatherer (EHG), Caucasus Hunter Gatherer (CHG), and Neolithic farmers (NEOL) ancestries in yellow, and mobility in orange. The position of the 95% credible intervals is used to identify the effect of the variables on the left, right and back burial sides. The posterior densities result from the models with all variables as predictors (baseline + ancestry + mobility). **B)** Delta WAIC scores have been calculated by subtracting the minimum WAIC score from all the scores. The dashed horizontal line represents the delta WAIC score for the baseline model (null model). The red points show the models with the lowest score (0), and therefore the ones that have the best fit. C) Delta DIC have been calculated using the same method as for delta WAIC scores. The red points show the models with the lowest delta DIC score (0).

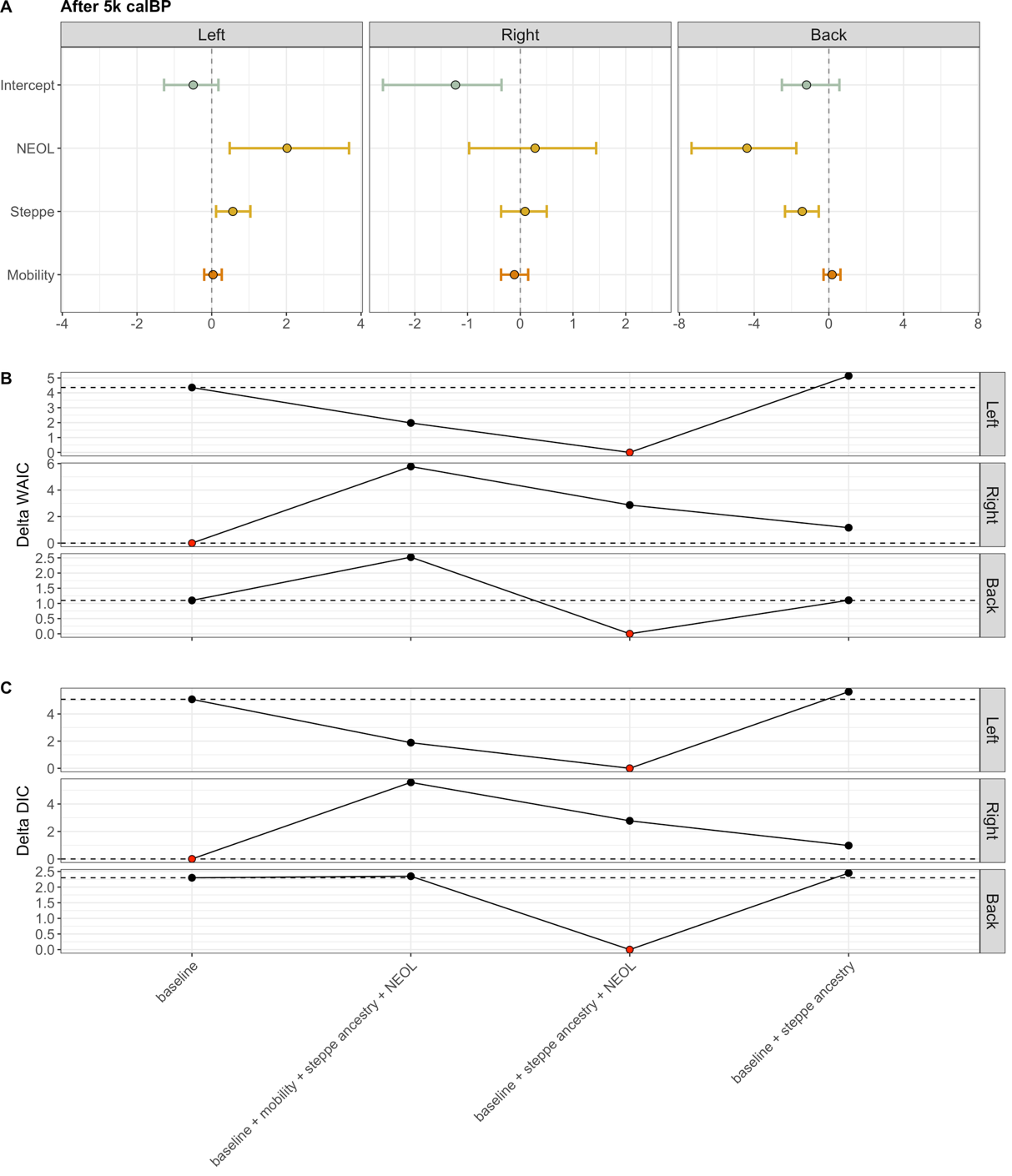

**Figure S11.** Spatiotemporal relationship of genetic ancestry and mobility with burial side after 5k calBP, when Steppe ancestry starts appearing in Western Europe. **A)** Posterior densities of the coefficients for the explanatory variables: the intercept in light blue, Eastern Hunter Gatherer (EHG), Caucasus Hunter Gatherer (CHG), and Neolithic farmers (NEOL) ancestries in yellow, and mobility in orange. The position of the 95% credible intervals is used to identify the effect of the variables on the left, right and back burial sides. The posterior densities result from the models with all variables as predictors (baseline + steppe ancestry + NEOL + mobility). **B)** Delta WAIC scores have been calculated by subtracting the minimum WAIC score from all the scores. The dashed horizontal line represents the delta WAIC score for the baseline model (null model). The red points show the models with the lowest score (0), and therefore the ones that have the best fit. C) Delta DIC have been calculated using the same method as for delta WAIC scores. The red points show the models with the lowest delta DIC score (0).

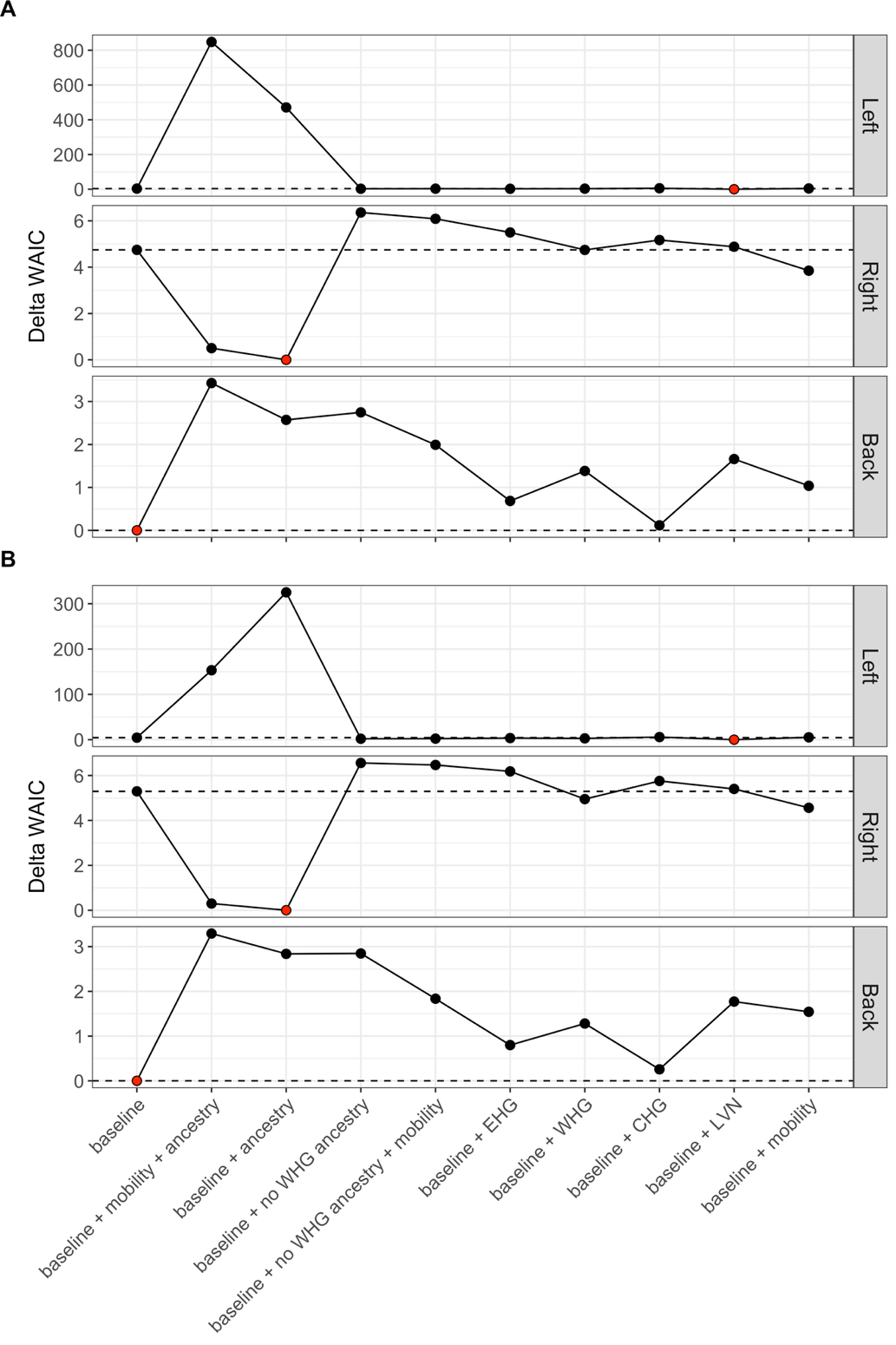

**Figure S12.** Delta WAIC (**A**) and delta DIC (**B**) scores for all model tested, including the ones with all ancestries (WHG, EHG, CHG, NEOL). Delta WAIC and delta DIC scores have been calculated by subtracting the minimum WAIC or DIC score from all the scores. The dashed horizontal line represents the delta WAIC/DIC score for the baseline model (null model). The red points show the models with the lowest score (0), and therefore the ones that have the best fit.

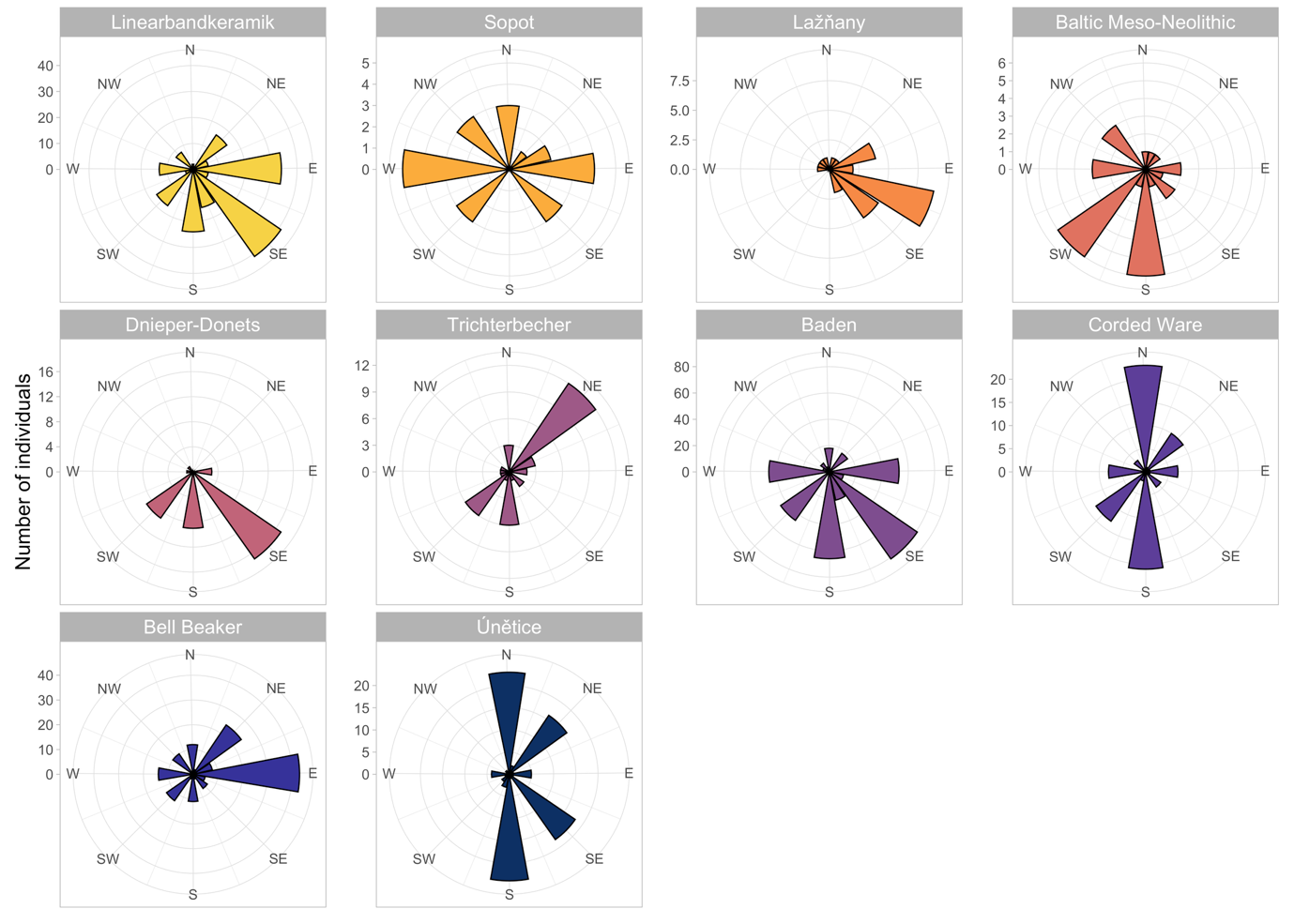

**Figure S13.** Burial orientation across cultures. Histograms show the number of individuals and their burial orientations, for graves belonging to the ten most represented cultures in our data, from oldest to youngest. Shown are all individuals grouped by culture, independently of ancestry information.

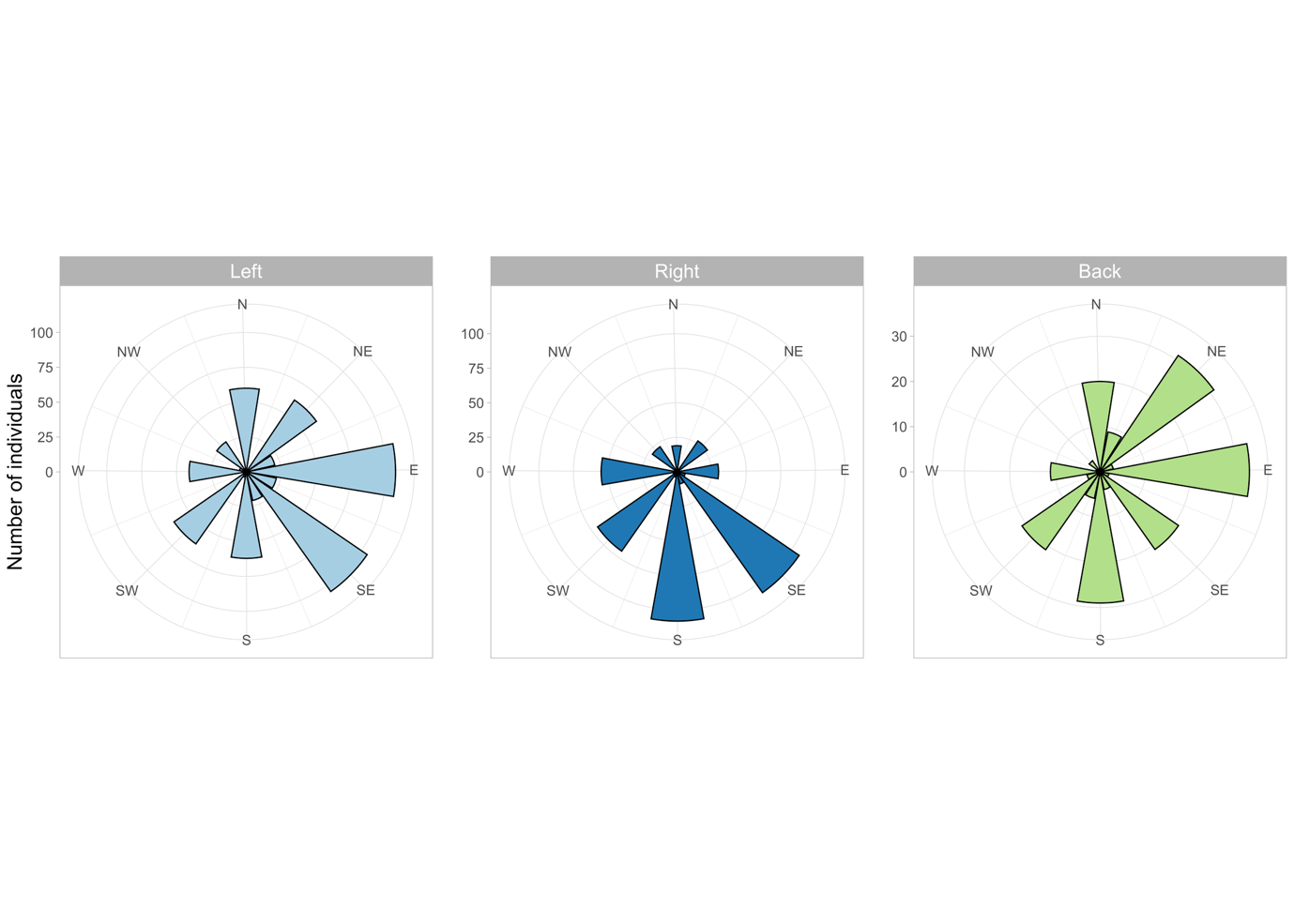

**Figure S14.** Burial orientation across burial sides. Histograms show the number of individuals and their burial orientations, grouped by the side they were buried (left, right, back).

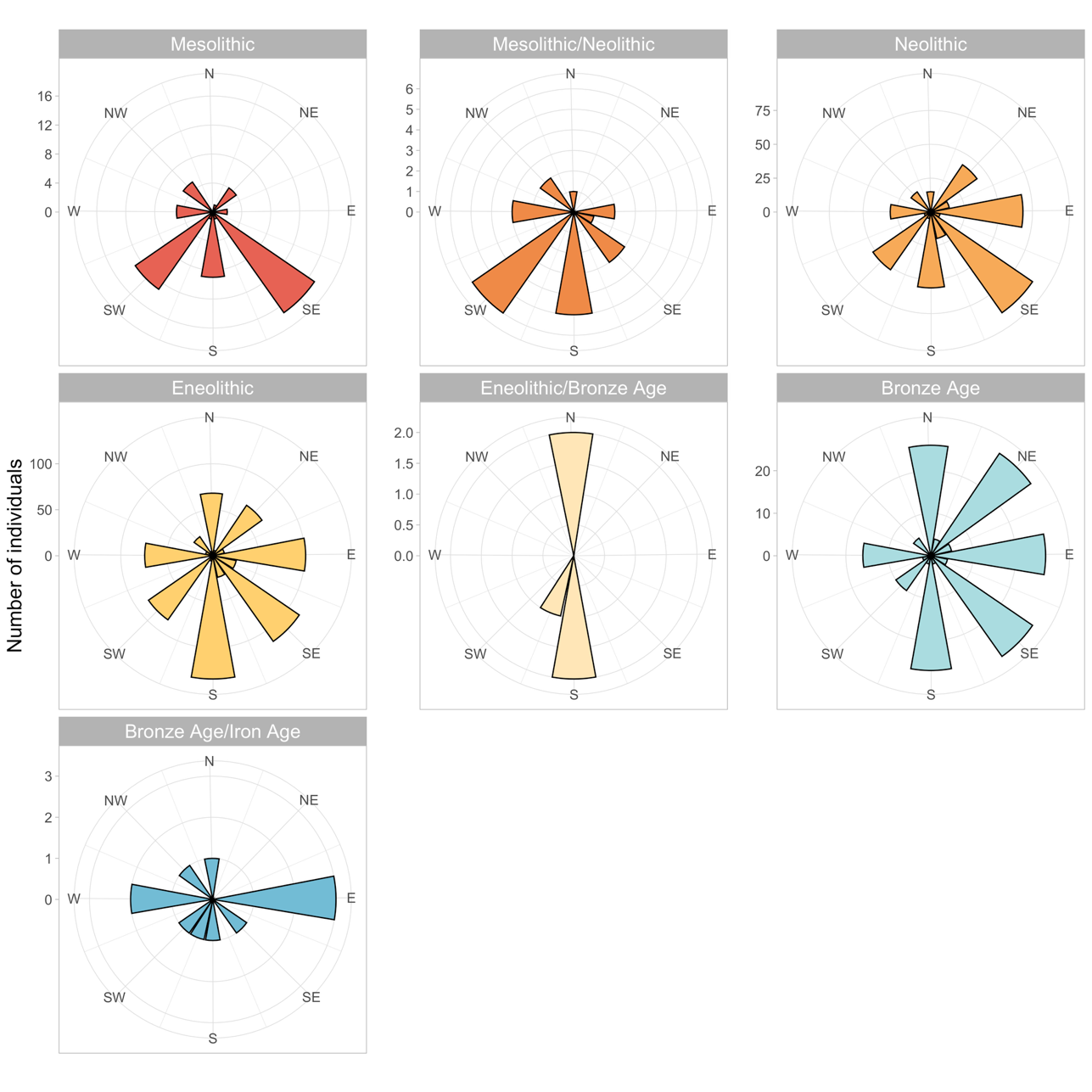

**Figure S15.** Burial orientation across archaeological period. Histograms show the number of individuals and their burial orientations for each period, from oldest to youngest.

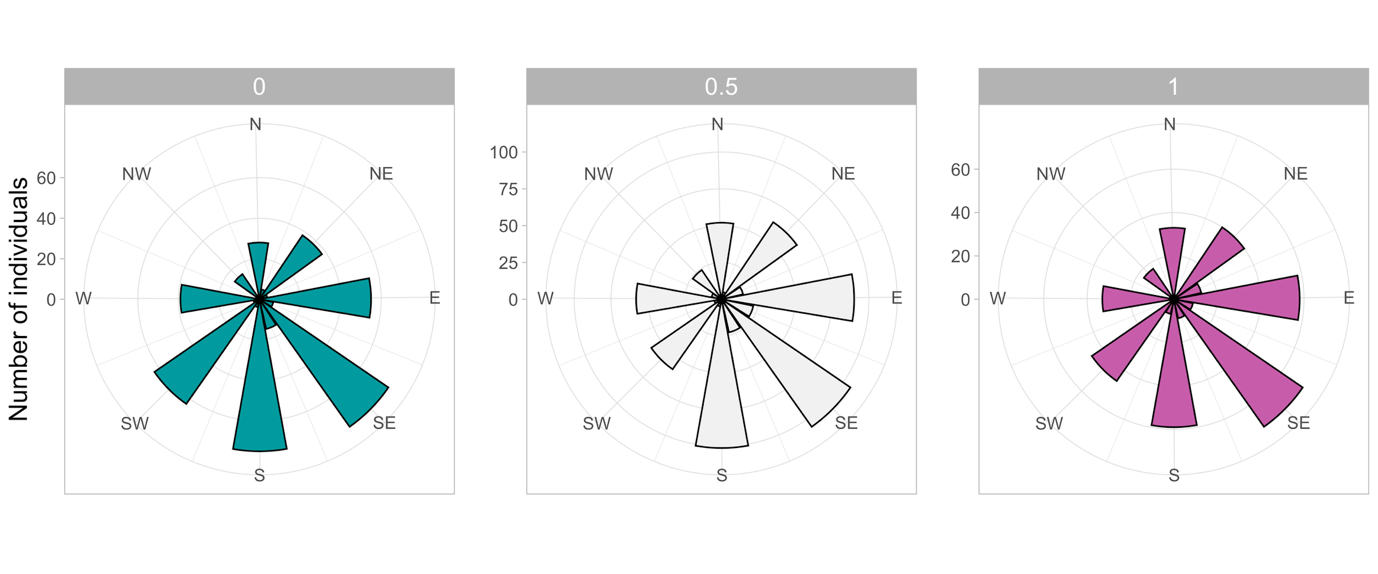
**Figure S16.** Burial orientation across sex of the individuals. Histograms show the number of individuals for each burial orientation and sex, assigned as a probability of being female based purely on the anthropoplogical evidence. A value of 1 indicates female with certainty, a value of 0 indicates male with certainty, and a value of 0.5 indicates an equal probability of male or female. In cases where the individual has not been osteometrically assessed, we assumed a prior of 0.5.

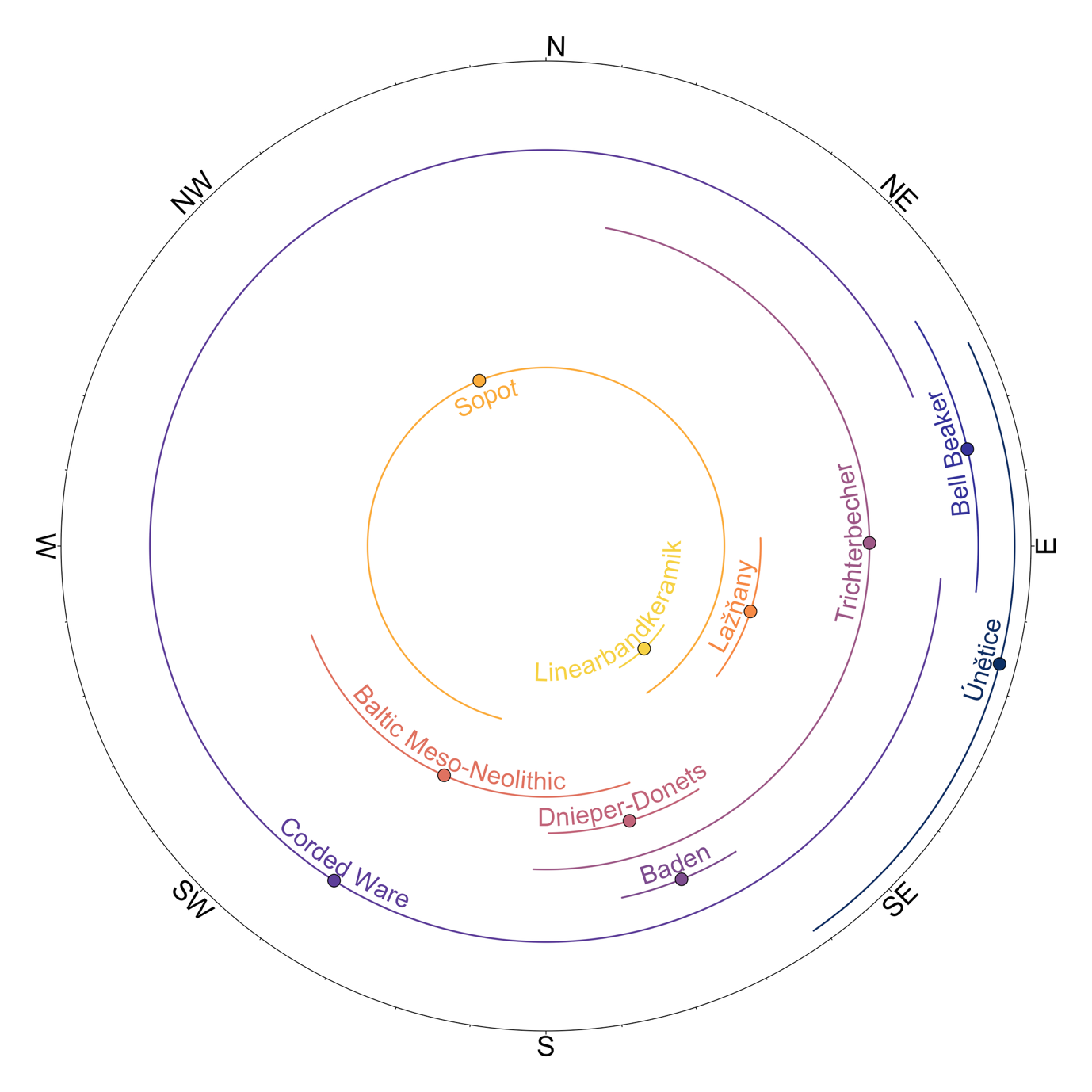

**Figure S17.** Results of the Bayesian circular regression testing for differences in burial orientation across the ten most represented cultures in our data. Shown are the posterior distributions of the circular means for each culture, with the points representing the mean of the distribution and the lines the upper and lower bounds. Cultures have statistically different circular means if their posterior distributions do not overlap. All individuals with culture information were included in the model.

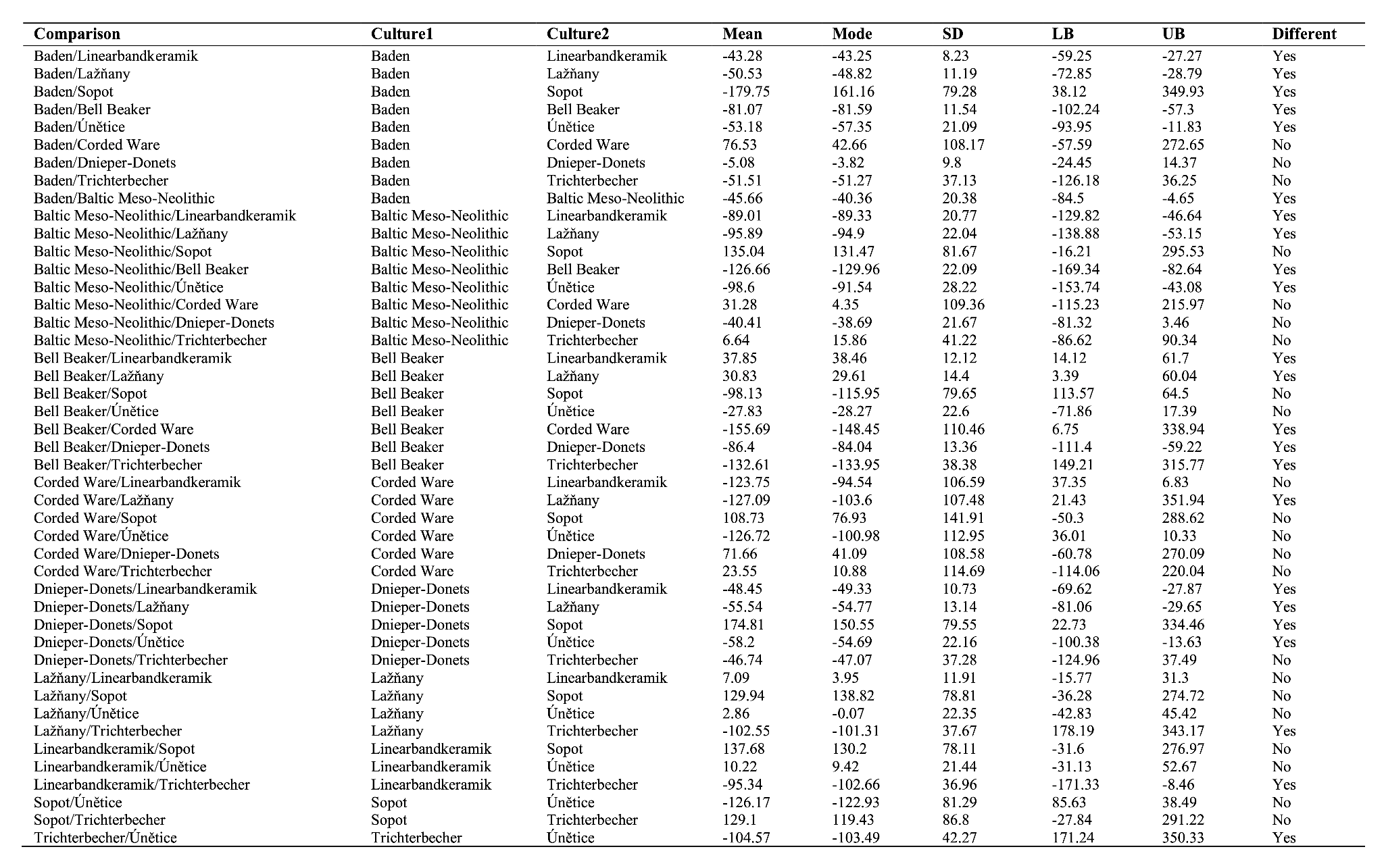
**Table S1.** Comparison of burial orientation across cultures. The table shows the results from a set of Bayesian circular regression models, indicating the difference (interpreted as angular distance along the clock) in the posterior distributions of orientations between pairs of cultures (**Culture1** and **Culture2**). Summary statistics are: the difference in circular mean (**Mean**), the difference in circular mode (**Mode**), the difference of the standard deviation (**SD**) and the differences between the lower (**LB**) and upper (**UB**) bounds of the 95% highest posterior density interval for the two cultures that are being compared. For example, the difference between circular means of Linearbandkeramik and Baden is:

$114.7245˚ \left( Linearbandkeramik \right)-158.0078˚ \left( Baden \right)= -43.28329$˚
